# Mapping Glycan Binding Profiles of the Gut Microbes using Liquid Glycan Array (LiGA)

**DOI:** 10.64898/2026.03.17.712435

**Authors:** Pravina Yadav, Chuanhao Peng, Mirat Sojitra, Shreyas Gupta, Ben Willing, Ratmir Derda

## Abstract

Glycan-microbe interactions are central to gut colonization and host-microbiota communication. Here, we apply a DNA-encoded Liquid Glycan Array (LiGA) to quantify interactions between live gut bacteria and multivalent natural or mirror glycans. LiGA comprises glycosylated M13 bacteriophage bearing silent DNA barcodes that encode glycan identity and density. Using LiGA, we profiled glycan binding across 16 *Limosilactobacillus reuteri* strains isolated from murine, porcine, poultry, and human hosts, and then extended the approach to profile glycan binding of taxonomically diverse bacteria from three phyla Bacillota, Bacteroidota, Pseudomonadota consisting of nine different species. Recent discussion of mirror-image microorganisms raise a question whether these microorganisms could interact with present-day life by engaging naturally chiral glycans. We demonstrated that this question can be assessed by testing the binding of mirror-image glycans to natural bacteria. Evaluation of enantiomers of common glycan revealed cross-chiral recognition by *Escherichia coli* and *L. reuteri*, indicating that these bacteria can used mirror-image glycans to engage for adhesion and potential colonization. By symmetry the same arguments extends to mirror microorganisms and glycans of naturally chirality. We established that LiGA enables efficient characterization of bacterial glycan binding and provides new insights into intestinal microbial ecology.

## Introduction

Dense coat of glycans is present on the surface of all cells across all kingdoms of life, highlighting their vital roles in communication between cells. Glycan recognition is linked to self versus non-self discrimination by immune systems, and recognition of glycans by glycan-binding proteins (GBPs) is the first line for detection of pathogenic microorganisms. To understand GBP-glycan interactions between host organisms and microbiome, technological advances are needed to quantify protein-glycan interactions with intact cells and ideally in true physiological context. Technologies such as lectin microarrays have been utilized for the analysis of glycosylation of bacteria^1^. Antibody and lectin microarrays have played an important role in illustrating bacterial glycosignatures, enabling differentiation among strains, and to evaluate the difference in the glycan structures due to changes in environmental conditions^2^. For observing host-bacteria glycan-glycan interactions, purified bacterial glycans have been used on the microarray printed host glycans^3^. Elegant LectinSeq technology that couples lectin-glycan recognition and metagenomic analysis made it possible to detect which glycan recognition events are important in native microbial communities^4^. This and other emerging examples that combine GBP-glycan recognition with Next-generation Sequencing (NGS) highlight the potential of NGS-powered technologies in dissecting glycan-GBP interactions at host-bacteria interface.

Contrast to the progress of lectin-array technologies, the potential of the glycan arrays for studying host-bacteria interactions has not been adequately explored. Among thousands of applications of glycan array to measure glycan-binding profile of purified GBPs, there are only scarce examples employing the same arrays to study the glycan binding profile of GBPs on the surface live bacteria^5^. Reasons for this imbalance might originate from technical challenges associated with analysis of live bacteria and solid glass arrays. Monovalent glycans tagged by DNA have been employed to overcome the challenges that display of glycans on solid surfaces might present^6^. However, monovalent glycans are not a suitable mimetic for a multivalent display of glycans on the surface of host cells, mucosa, and other biological interfaces with which bacteria may interact. We have previously disclosed a strategy for DNA-encoding of a controlled multivalent display of glycans on the surface of M13 bacteriophage^7, 8^. The Liquid Glycan Array (LiGA), a mixture of such DNA-encoded multivalent glycosylated constructs, is ideally suited for measurement of binding profile of GBPs on the surface of live mammalian cells *in vitro* and *in vivo*^7-10^. Unlike the “glycophage display” system that employs the biosynthesis of glycans in bacteria^11^, LiGA technology offers multivalent presentation of any synthetic glycans at desired density. Specifically, LiGA employs chemical ligation of glycans to a subset of ∼2700 copies of major coat protein pVIII to produce a multivalent display of ∼150-1500 copies of glycans, and LiGA retains most of the benefits of other M13 phage-displayed libraries, importantly, robust protection of the DNA message inside the virion, which has allowed successful profiling of ligand-receptor interactions in organs in live animals^8, 12^ and in humans^13^. The solution-phase format of LiGA assay, in principle, makes it amenable to screen glycan binding of intact bacterial cells and communities. In this manuscript, we provide the first comprehensive assessment of measuring glycan binding of live prokaryotes with LiGA and highlight technological plausibility for *in vitro* profiling of GBPs on the surface of bacteria, and in the future even in microbial communities *in vivo*.

The gut microbes exhibit a phenomenal strain level diversity^14^. Despite significant progress in understanding the genomic basis of this diversity^15^. The study of functional basis of this diversity is still in nascent stage^16^. The ability of gut microbes to forge competitive or cooperative relationships with one another is influenced by their physical proximity^17^. Glycan mediated adhesion is important for bacteria to increase their access to nutrients in diverse ecosystems such as adhesion of marine microbes to chitin^18^ and adhesion of cellulolytic species to cellulose^19^. Moreover, pathogenic microbes adhere to host cells to enable access to nutrients^20^; in many cases, adhesins bind to specific host glycan structures^21^. Besides pathogenic microbes, commensals such as *Lactobacilli* and *Bacteroides* interact with host glycans using mucus-binding proteins (MUBs) and surface glycan binding proteins (SGBPs) respectively. Gut microbes not only enhance the metabolic ability of the host through the nutrient provision but also exclude pathogens and help in the host immune system^22^. The taxonomic profile of the vertebrate microbiota is largely host-specific^22, 23^ and in some cases, congruent with the evolution of host species. Little is known about the mechanisms by which gut microbes can evolve stable associations with their hosts that would allow for reciprocal evolutionary interactions between bacterial lineages and host^24^. The gram-positive bacterium *Limosilactobacillus reuteri* has been used as a model organism to study the evolutionary strategy of vertebrate gut symbionts because this organism colonizes the gastrointestinal (GI) tract of vertebrates as diverse as humans, rodents, pigs, and chickens. In rodents, pigs and chickens, it is one of the dominant species in the GI tract and forms the biofilm-like associations with the stratified squamous epithelial lining of the proximal regions of the digestive tract^25^. It is observed that strains of *L. reuteri* from global sources comprised distinct phylogenetic clusters that can be detected with the Multilocus Sequence Analysis (MLSA) and Amplified Fragment Length Polymorphism (AFLP), and these clades show significant association with host origin^26^. Besides these genotypic patterns, an adaptive evolutionary process is also demonstrated by phenotypic characteristics of *L. reuteri* strains in terms of the ecological performance in the gut and adhesion to epithelial cells^27^. Genomic approaches in combination with experiments in animal models offer mechanistic insight into the evolution and ecology of microbial symbionts of vertebrates. A microarray analysis of 57 *L. reuteri* strains revealed specific gene combination in host adapted lineages of *L. reuteri*^24^. Several of these surface proteins are predicted to be involved in epithelial adhesion, mucin binding, and extracellular matrix binding^24^. Although adhesion of these surface proteins to epithelial cells is known to be mediated by glycans, direct binding of intact *L. reuteri* with specific glycans remains to be demonstrated. Here, we examined the role of glycan binding profiles of *L. reuteri* in the host specificity and explored glycan binding of taxonomically diverse bacteria from three different phyla Bacillota, Bacteroidota, Pseudomonadota consisting of 9 different species (*L. reuteri, Escherichia coli, Bacteroides dorei, Bacteroides thetaiotamicron, Bacteroides fragilis, Bacteroides vulgatus, Lactobacillus mucosae, Citrobacter fruendii*, and *Clostridium ramosum*).

All known life is homochiral. Advances in synthetic biology have enabled the creation of functional component of putative mirror life. The biomolecules, the biochemical components of the Central Dogma with chirality opposite to that of natural ones, can give rise to mirror-image *in vitro* translation system, which could offer benefits such as the production of proteolytically resistant biomedicines. However, a recent assessment of the potential risks associated with mirror-image life forms^28^, together with a related technical report^29^, has brought an important issue to the attention of both the scientific community and the public. Major concerns were raised by the authors, who argued that many aspects of the host response to mirror bacteria could be deficient. A follow-up commentary^30^ further emphasized the critical importance of glycoconjugates in evaluating the risks posed by putative mirror-image life. Glycans, in addition to proteins and nucleic acids, must be considered to reach an accurate assessment of our preparedness for the arrival of mirror-image organisms. Enantiomeric pairs of monosaccharides have already been identified in the glycocalyx of many extant organisms^31^. We have recently reported cross-chiral recognition of enantiomeric glycans by naturally occurring glycan-binding proteins^32^. We further reasoned that the potential colonization of mirror bacteria in hosts bearing naturally chiral glycans can be examined by measuring the adhesion of native bacteria to multivalent displays of mirror glycans. Aside from classical analysis of chemotaxis towards enantiomers of common sugars, the extent to which bacteria can adhere to glycan enantiomers has not been systematically studied^33^. In this manuscript, we use LiGA to uncover the binding of enantiomers of common monosaccharide to intact *E. coli* and *L. reuteri*, which represented common entero-bacteria circulating in global population and constantly interacting with human glycome.

## Results

### Validation of the LiGA glycan-binding assay with intact wild type and ΔFimH *E. coli*

To establish the efficiency of “liquid format” of binding between glycosylated bacteriophages and bacteria, we measured binding of LiGA-ED, comprising 81 glycans that represent most host glycan structures present on the gut epithelium and mucus layer (**Table S1**), to wild-type *E. coli* BW25113 and a fimH-deficient (ΔFimH) mutant strain. FimH is a mannose-binding adhesin expressed by wild-type *E. coli*.

To describe glycan identity and display density, each glycan-phage conjugate is denoted as “glycan-[density]”, where the bracketed number indicates the average number of glycans displayed per phage particle, as quantified by mass spectrometry. The Differential enrichment (DE) analysis confirmed strong enrichment of densely mannosylated bacteriophages (αMan-[840], (Man)3-[1300], and (Man)3-[1730]) following incubation of LiGA with wild-type *E. coli* BW25113, whereas the ΔFimH strain exhibited markedly reduced enrichment of these same mannosylated phages (**Figure 1D–E**). Mannose binding events were also being observed in DE analysis that employed the wild-type *E. coli* BW25113 as ‘test’ and the ΔFimH strain as ‘control’ (**Figure 1F**). The combined observations confirmed successful application of LiGA to detect specific glycan-binding events with receptors on the surface of intact bacteria. Direct analysis of binding, exemplified as DE comparison vs. the input composition, and differential analysis of two strains, comparison of FimH^+^ and FimH^-^ *E. coli*, were viable strategies for pan-surface interaction with glycans or characterization of receptor-specific events. LiGA analysis of GBPs on bacterial cells is conceptually and technologically follows already reported analyses of GBPs on the surface of mammalian cells^9^. Major technical differences were in isolation of LiGA-borne DNA barcodes associated with bacterial cells: neither boiling^10^ nor proteinase/RNAse treatments^9^ of bacteria pellets were successful, whereas a standard miniprep isolation of plasmids from the bacteria followed by PCR of specific LiGA plasmid proved to be the most effective (**Figure 1B**).

**Figure 1.**
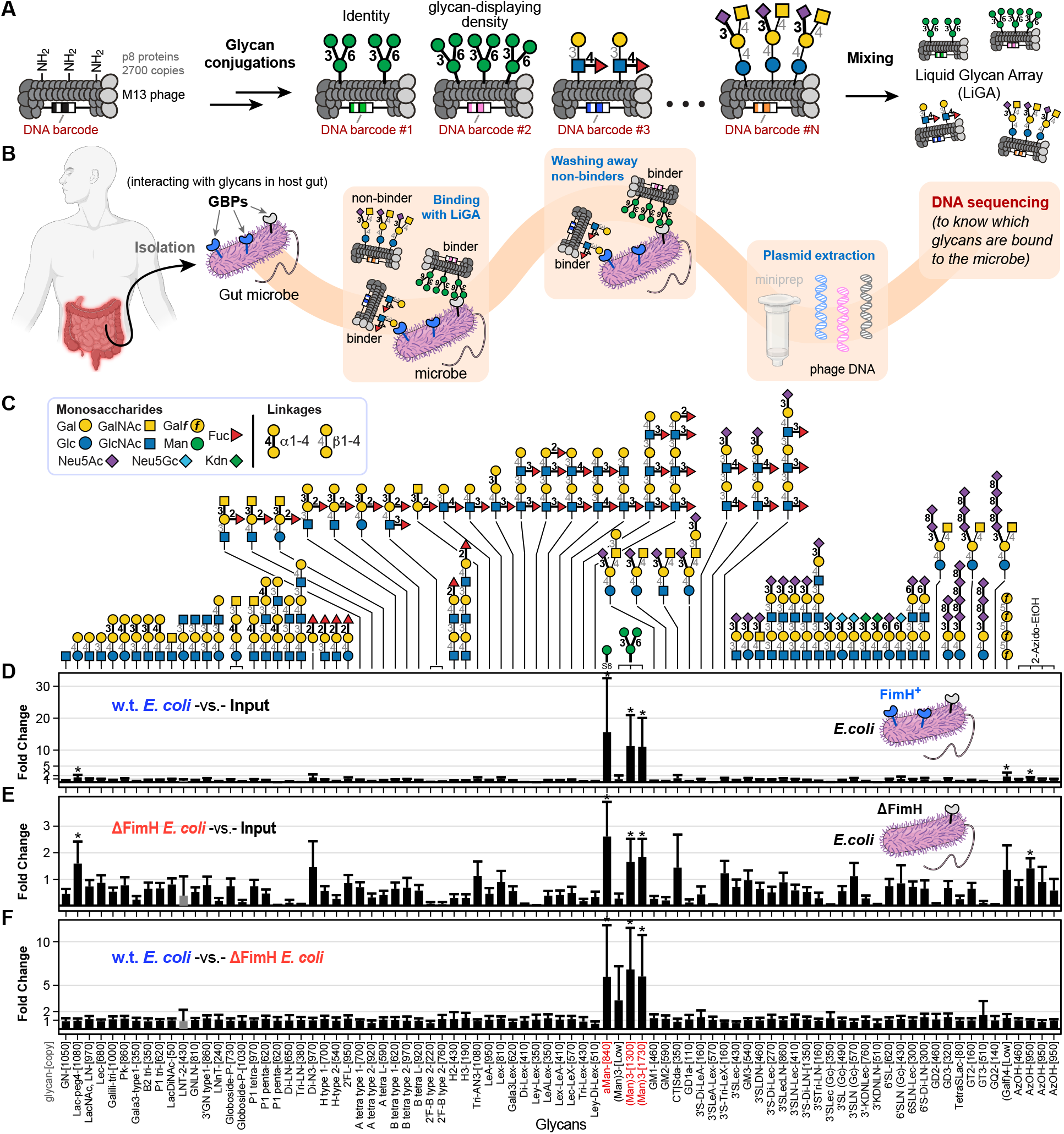
Profiling glycan binding of *E. coli* using LiGA. (A) Generation of LiGA. (B) Workflow of profiling glycan binding of gut microbes using LiGA. (C) Structure of glycans in the LiGA library. d–f, Glycan binding profiles of wild-type *E. coli* BW25113 versus naïve input library (D; test, n = 12; control, n = 4), ΔFimH mutant *E. coli* BW25113 versus naïve input library (E; test, n = 6; control, n = 4), and wild-type versus ΔFimH mutant *E. coli* BW25113 (F). Fold change (FC) was calculated by edgeR DE analysis using the negative binomial model, TMM normalization and BH correction for False-Discovery Rate (FDR). All respective n values are independent biological replicates, and FC + s.d. are reported with asterisks indicating FDR < 0.05. GBPs: glycan-binding proteins; FimH: type 1 fimbriae D-mannose specific adhesin. Panel (B) was created with BioRender.com.

### Glycan binding of the *L. reuteri* strains from different hosts does not explain host specificity

To determine whether *L. reuteri* strains from different hosts exhibit distinct glycan-binding profiles that could account for host specificity, we measured glycan-binding profiles of 16 *L. reuteri* strains isolated from poultry, porcine, rodents, and human using LiGA (**Figure 2** and **Figure S1–S4**). The glycan binding of those isolates was summarized as a heatmap based on hierarchical clustering of k-means of the Euclidean distance (**Figure 2A**) and the structure of enriched glycans was shown in **Figure 2B**. Contrary to our expectation, the strains of *L. reuteri* originating from the same host lineage did not cluster together based on their glycan-binding profiles. Although each strain displayed a unique glycan-binding profile, there are noticeable similarities of these isolates. All strains bound to at least one of the three mannose-containing motifs presented in the LiGA, and all strains showed binding to glycans terminating in galactose. Eleven out of the sixteen isolates demonstrated significant binding to Lac-peg4-[1080] (lactose), and seven out of the sixteen isolates recognized Di-N3-[970] (O blood group disaccharide), with *L. reuteri* JCM1081 exhibiting the strongest to both glycans. Although the most *L. reuteri* strains we tested shared substantial glycan-binding similarities, certain isolates exhibited unique preferences. Specifically, only *L. reuteri* limo showed binding to Tri-AN3-[1080] (A blood group trisaccharide), and only *L. reuteri* AP3 bound to 3′S-Di-Lec-[270] (**Figure 2**).

**Figure 2.**
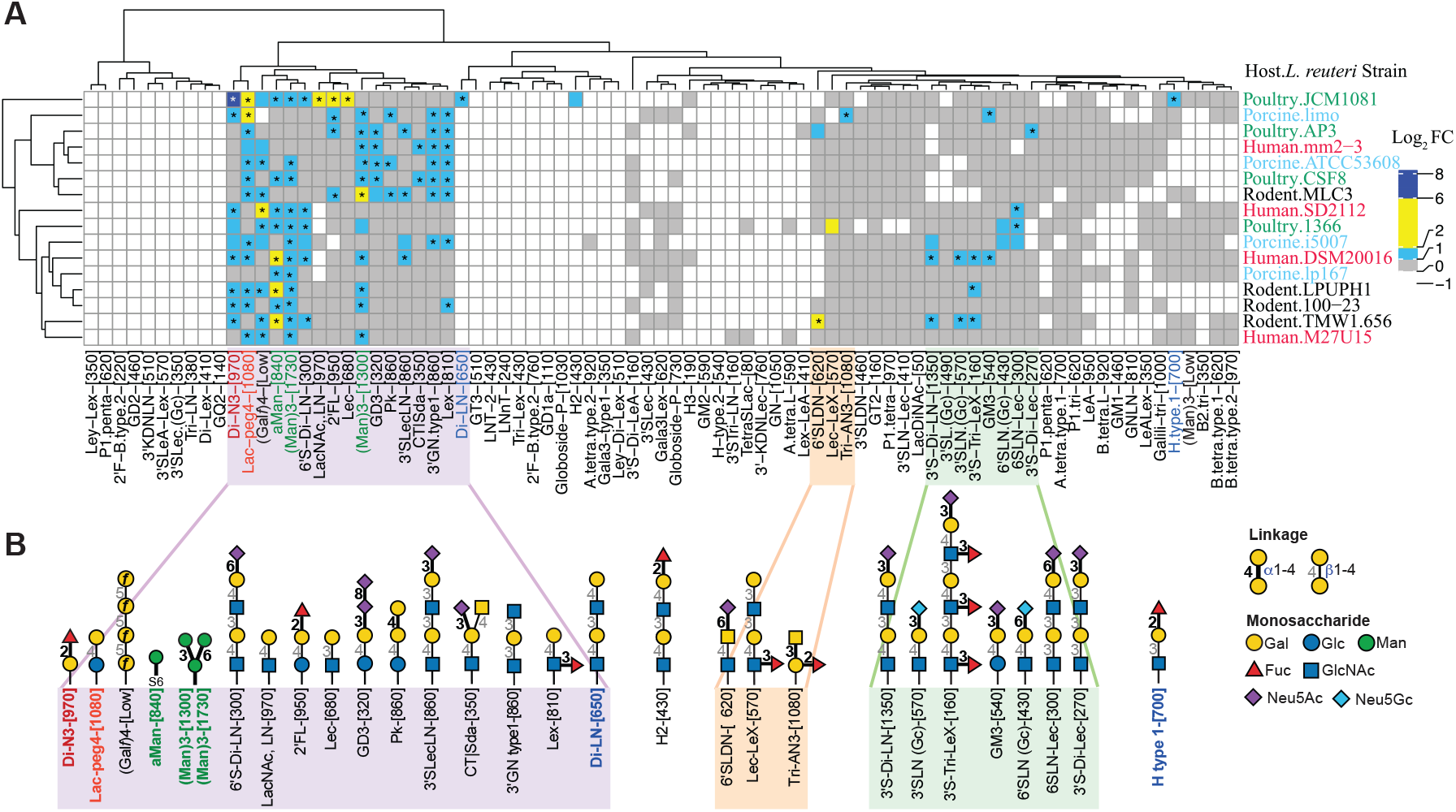
Heatmap of glycan binding of 16 *L. reuteri* strains using the hierarchical clustering of k-means of the Euclidean distance. Structure of enriched glycans were shown in panel (B). * Represents FC ≥ 2, FDR ≤ 0.05, n = 7. Log_2_FC was calculated by edgeR DE analysis using the negative binomial model, TMM normalization and BH correction for FDR. Isolates from different hosts are marked in distinct colors. Heatmap was drawn using pheatmap package in R.

We further pulled out LiGA binding data of four poultry *L. reuteri* strains—JCM1081, CSF8, 1366, and AP3 to examine whether strain-specific glycan-binding differences exist among isolates from the same host (**Figure S1**). We found that seven glycans were significantly enriched with *L. reuteri* JCM1081. Among these, Di-N3-[970] and Lac-peg4-[1080] exhibited markedly higher binding to JCM1081 than to the other isolates, making JCM1081 an outlier in the hierarchical clustering of all *L. reuteri* strains.

### Glycan binding of a poultry-isolate *L. reuteri* JCM1081 identifies it as a unique strain

Among all *L. reuteri* strains tested, *L. reuteri* JCM1081 exhibited the strongest binding to multiple glycans terminating in galactose (Lac-peg4-[1080], LacNAc,LN-[970], and Lec-[680]), fucose (Di-N3-[970] and 2’FL-[950]), and mannose (αMan-[840], (Man)3-[1300], and (Man)3-[1730]) (**Figure 2** and **Figure S1A**). This pronounced glycan-binding suggests a high colonization potential in environments enriched in these glycans. Consistent with this interpretation, *L. reuteri* JCM1081 has been reported to be among the most adhesive strains isolated from poultry, exhibiting strong binding to mucin and human enterocyte-like HT-29 cells *in vitro*^34^. These observations motivated us to further investigate whether *L. reuteri* JCM1081 can also interact with mirror-image glycans. Addressing this question allows us to assess the potential colonization capacity of a hypothetical mirror *L. reuteri* in a host environment dominated by naturally chiral glycans.

### Glycan binding of taxonomically diverse gut bacterial isolates shows strain-level variations

Profiling glycan binding of the 16 strains of *L. reuteri* with LiGA has led us observed the strain level glycan-binding variations. To test if it holds true that strain-level glycan-binding variations also exist between other gut bacterial strains isolated from different hosts, we measured glycan binding of taxonomically diverse bacteria from three phyla Firmicutes/Bacillota, Bacteroidota/Bacteroidetes, Proteobacteria/Pseudomonadota (consisting of 9 different species *L. reuteri, E. coli, B. dorei, B. thetaiotamicron, B. fragilis, B. vulgatus, L. mucosae, C. fruendii*, and *C. ramosum*) (**Figure 3** and **Figure S1–S6**).

**Figure 3.**
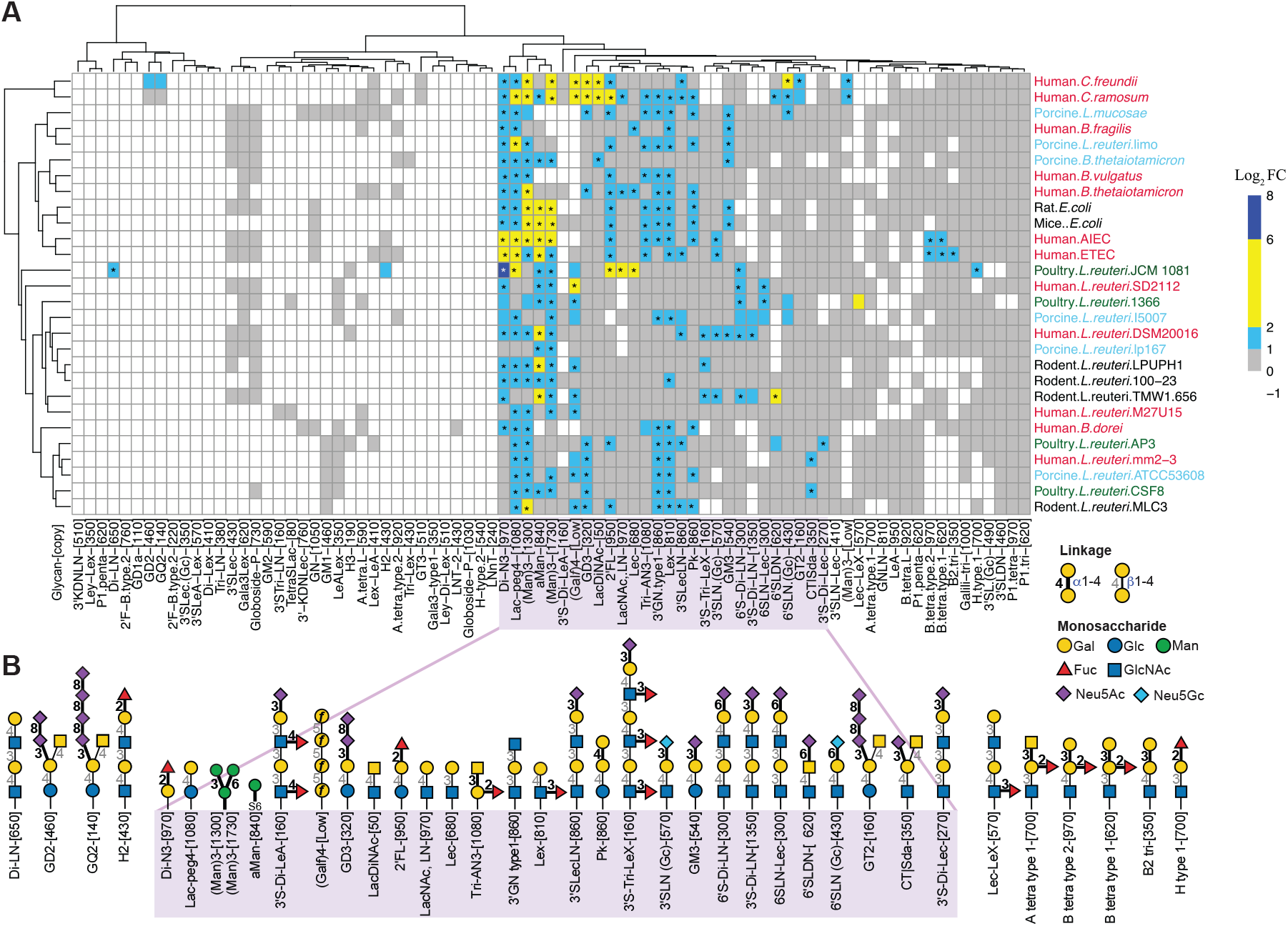
Summary of glycan binding of different isolates from three phyla, five genera and nine different species. Bacteria are from there phyla, Proteobacteria (Pseudomonadota), Bacteroidetes (Bacteroidota), and Firmicutes (Bacillota). Heatmap is drawn using the hierarchical clustering of k-means of the Euclidean distance with complete linkage clustering method. (* represents FC ≥ 2, FDR ≤ 0.05, n = 7). Structures of enriched glycans were shown in (B). Log_2_FC was calculated by edgeR DE analysis using the negative binomial model, TMM normalization and BH correction for FDR. Heatmap was drawn using pheatmap package in R. Isolates from different hosts are marked in distinct colors.

We observed that hierarchical clustering (using complete-linkage method) according to the glycan binding profiles of all bacteria showed three primary clades in the dendrogram (**Figure 3**). Clade 1 comprised *C. freundii* and *C. ramosum*. Clade 2 was subdivided into three subclades: first included *L. mucosae, B. fragilis*, and *L. reuteri* limo; the second contained *B. thetaiotamicron* isolated from pig and human together with *B. vulgatus*; and the third consisted of all *E. coli* strains. Clade 3 comprised 15 *L. reuteri* strains along with *B. dorei*. The *C. freundii* and *C. ramosum* within the Clade 1 exhibited strong binding to Lac-peg4-[1080], (Man)3, (Gal*f*)4-[Low], GD3-[320] (Neu5Acα2-8Neu5Acα2-3Galβ1-4Glcβ-Sp), LacDiNAc-[50] (GalNAcβ1-4GlcNAcβ-Sp), 2’FL-[950], and 6’SLN(Gc)-[430] (Neu5Gcα2-6Galβ1-4GlcNAcβ-Sp) (**Figure 3** and **Figure S6**). These glycans are not bound by other bacteria except the porcine *B. thetaiotamicron* binding to LacDiNAc and *L. mucosae* binding to 6’SLN(Gc). These different glycan binding features of *C. freundii* and *C. ramosum* led the clustering of these two in a separate clade.

The third subclade within the Clade 2 comprised all *E. coli* strains due to their similar binding to Di-N3-[970] (H blood group disaccharide), 2’FL-[950] (H blood group type 6), Tri-AN3-[1080] (A blood group trisaccharide), Le^x^-[810] (Galβ1-4[Fucα1-3]GlcNAcβ-Sp), Lac-peg4-[1080], and mannose-containing glycans (αMan-[840], (Man)3-[1300], and (Man)3-[1730]). However, there are some differences in glycan binding of *E. coli* commensals and *E. coli* pathogenic strains as 3’SLN(Gc)-[570] (Neu5Gcα2-3Galβ1-4GlcNAcβ-Sp), B tetra type 2-[970] (Galα1-3[Fuca1-2]Galβ1-4GlcNAcβ-Sp), and B tetra type 1-[620] (Galα1-3[Fucα1-2]Galβ1-3GlcNAcβ-Sp) are bound by the pathogenetic adhesive invasive *E. coli* (AIEC) and enterotoxigenic *E. coli* (ETEC) but not by commensal strains. GM3-[540] (Neu5Acα2-3Galβ1-4Glcβ-Sp) was bound by commensals but not by the pathogenic strains. Two glycans B2-tri-[350] (Galα1-3Galβ1-4GlcNAcβ-Sp) and 3’SLecLN-[860] (Neu5Acα2-3Galβ1-3GlcNAcβ1-3Galβ1-4GlcNAcβ-Sp) were only bound by ETEC. Subclade 2 and 3 of Clade 2 has four Bacteroidetes and two Firmicutes due to similar glycan binding (**Figure 3**).

The heatmap (**Figure 3**) illustrates the similarity in glycan binding of bacteria from similar taxonomic groups shown by cluster of pathogenic and commensal *E. coli* strains (third subclade of the Clade 2) and 15 strains of *L. reuteri* (Clade 3). However, there are a few exceptions, such as *C. freundii and C. ramosum* cluster together despite being taxonomically distant.

Glycan-binding preferences likely reflect bacterial adaptation to the glycan composition of the gut environment. Mucin O-glycans are rich in Gal, GalNAc, GlcNAc, and Fuc, but not in mannose^35,36^. However, it has been reported that mannose is abundant in epithelial layer^37^ and dietary glycans^38^. Thus, the ability of isolates in our study to bind both terminal galactose and mannose suggests they interact with both mucin-derived and epithelial surface glycans in the gut.

All *Bacteroides* strains bound to galactose, mannose and fucose containing glycans while only *B. thetaiotamicron* and *B. fragilis* bound to sialic acid containing glycans GD3 (Neu5Acα2-8Neu5Acα2-3Galβ1-4Glcβ-Sp) and GM3 (Neu5Acα2-3Galβ1-4Glcβ-Sp) (**Figure 3** and **Figure S5**). In *B. thetaiotamicron*, SusD-like protein BT1043 (outer membrane protein) has been associated with O-glycan utilization of host mucin^39^. A SusD like protein NanU, a SusD family protein from *B. fragilis* has also shown high binding affinity to sialic acid^40^. *B. vulgatus* and *B. dorei* did not bind to any sialic acid containing glycan. Possible explanation is that sialic acid is mainly present in the host epithelium and mucus layer glycans and rarely in the dietary glycans, thus these dietary glycan recognizing/metabolizing bacteria have evolved to interact mildly with host glycans containing sialic acid^41^. Mechanism of interaction of these bacteria with host has not been studied, therefore, further study will be required to show that *B. dorei* and *B. vulgatus* do not bind to sialic acid. *Bacteroides* binding to the host mannose and galactose containing (Lac-peg4, Di-N3 etc.) structures in glycan array is speculated to represent the *Bacteroides* binding to dietary mannan (Man)_n_ and galactan (Gal)_n_^17^. It is speculated that the glycan-binding that we observed may also indicate that binding of members of *Bacteroides* is mediated by cell surface lectins; for example, lectins incorporated into pilli that extend beyond polysaccharide capsule^42^, and surface anchored CAZymes/GHs (with carbohydrate binding modules).

*C. freundii* and *C. ramosum* showed strong binding to LacdiNAc (GalNAcβ1-4GlcNAcβ-Sp), which is especially expressed in MUC5AC gastric mucins^43^, and LacdiNAc has been associated with the tissue specificity of the *H. pylori* in gastric tissues. Tissue tropism of *H. pylori* has been suggested to be the result of strong acidic conditions of the stomach^44^. BabA and SabA adhesins of *H. pylori* have been characterized to bind Le^b^ and sialylated Le^a^ or Le^x^ respectively. However, functional binding of these adhesins cannot explain tissue tropism as these glycan motifs are not restricted to gastric mucosa and are present on other parts of GI tract. Conversely, LabA (LacdiNAc-binding adhesin) of *H. pylori* explains its restricted specificity to the gastric mucous surface in GI tract as LacdiNAc co-localizes with MUC5AC of gastric mucosa^43^. LacdiNAc has also been associated with diverse micro-environments (other than GI tract) such as hepatic granuloma tissues^45^ and other diseased tissues^45, 46^. The human *C. freundii* and *C. ramosum* are known for systemic infection capability. *Erysipelatoclostridium ramosum* (also known as *C. ramosum*) is associated with disease in humans^47^ and *C. freundii* is bacteremic/speticemic, many organs and tissues are affected besides the gut^48^. It can be speculated that systemic infection ability of these bacteria is contributed by their LacdiNAc-binding adhesins. Further validation of adhesin is warranted as with exception of binding of *H. pylori* to LacdiNAc, direct binding of this glycan motif with other microbes has not been shown. In our glycan array analysis of porcine *B. thetaiotamicron*, its binding to LacdiNAc is unanticipated as this bacterium is neither seen in gastric mucosa nor colonizes other tissues except GI tract; therefore, further investigation of its LacdiNAc binding is warranted.

### Conserved glycan-binding profiles among diverse *E. coli* strains

Both human-derived pathogenetic *E. coli* strains, AIEC and ETEC, exhibited binding to 13 glycan structures, while rat- and mouse-derived *E. coli* strains showed binding to 11 glycans (**Figure 3**). All strains exhibited binding to Lac-peg4, Di-N3, 2’FL, Tri-AN3, and Le^x^, with particularly strong affinity for mannose-containing glycans at high displaying densities (**Figure 4A–D)**.

**Figure 4.**
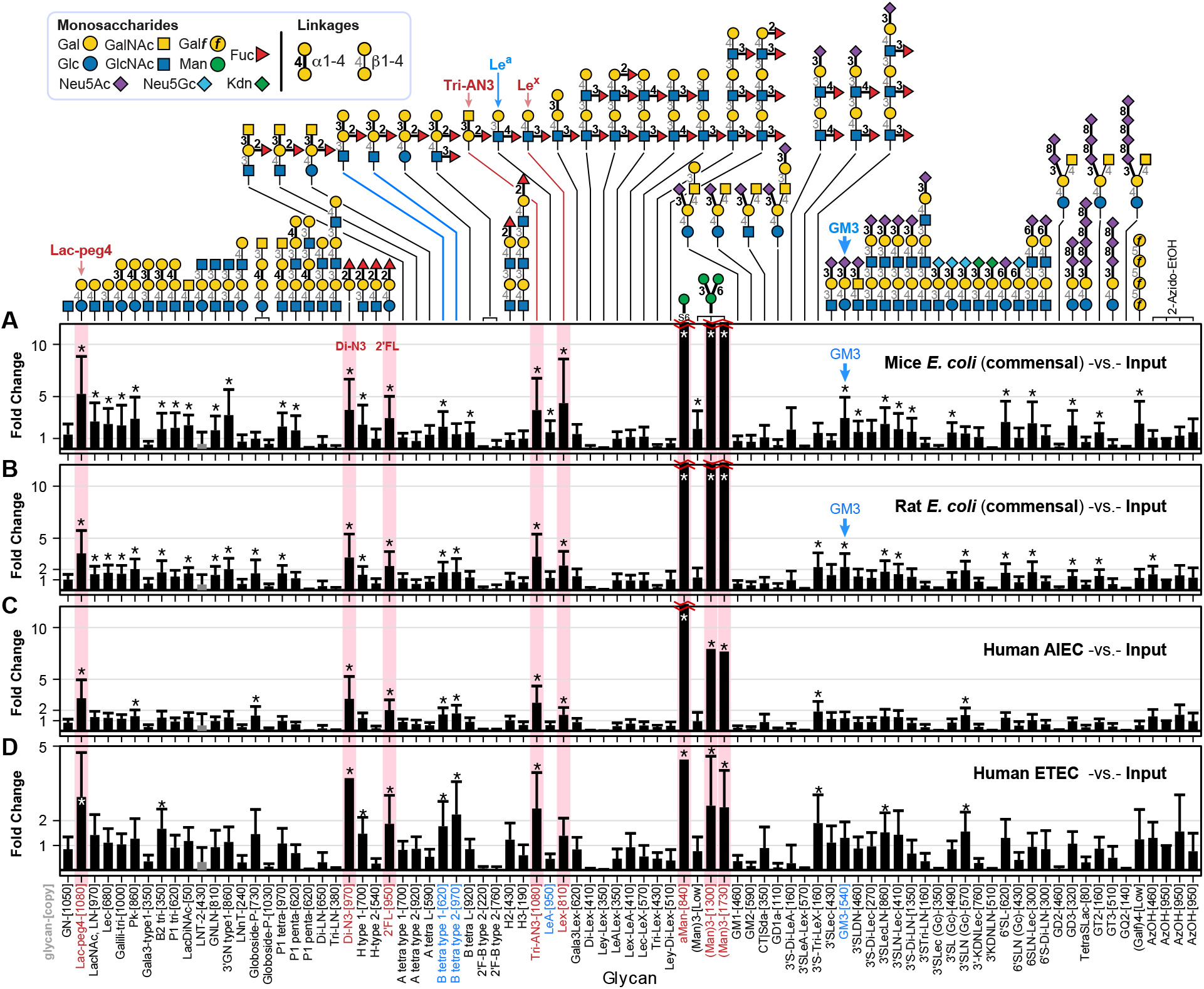
Glycan binding profiles of the *E. coli* strains. (A) Mice commensal *E. coli*, (B) Rat *E. coli*, (C) Adherent invasive *E. coli* (*AIEC*), and (D) Enterotoxigenic *E. coli* (*ETEC*). Fold change was calculated by Bioconductor edgeR DE analysis using the negative binomial model, TMM normalization and BH correction for FDR. Error bars represent s.d. propagated from the variance of the TMM-normalized sequencing data. * represents FDR ≤ 0.05, n = 7.

All four gut *E. coli* strains showed strong binding to mannose residues at high displaying densities, which supports the presence of FimH protein on the surface of *E. coli*^49^. In LiGA analysis, *E. coli* showed binding to Man-, Gal-, Fuc- and Sia-containing glycans, which correlated to the result of a recent study that probed bacterial fimbria-glycan interactions using *E. coli*^50^. Docking analysis showed *E. coli* Top10-CFA/I binding to Le^a^ (Galβ1-3[Fucα1-4]GlcNAcβ-Sp)^51^; however, our glycan array analysis did not show its binding to Le^a^-[950]. Instead, a weak binding to Le^x^-[810] (Galβ1-4[Fucα1-3]GlcNAcβ-Sp) was observed (**Figure 4A–D**). It might be result of strain level differences in glycan binding. Binding of AIEC and ETEC to the B tetra type 1 (Galα1-3[Fucα1-2]Galβ1-3GlcNAcβ-Sp) (Blood group B type 1) and B tetra type 2 (Galα1-3[Fucα1-2]Galβ1-4GlcNAcβ-Sp) (Blood group B type 2) had been previously shown in the FedF adhesin of ETEC^52^. In our study, ganglioside GM3 bound commensal *E. coli* strains and in a previous study, *E. coli* K99 fimbriae have shown binding to GM3 in intestinal tissues^53^.

### Cross-chiral recognition between enantiomeric glycans and live bacteria

Glycans are among the most abundant and prevalent biomolecules in nature and densely decorate the surface of all cells. To mediate adhesion and potentially colonization, microbes have evolved diverse glycan-binding proteins (GBPs) that recognize host cell glycans. We hypothesized that a putative mirror-image microorganism could similarly employ mirror-image GBPs to engage the host glycocalyx, thereby enabling adhesion and colonization. To profile glycan binding by such microorganisms that do not yet exist in nature, we envisioned that measuring the binding of glycan enantiomers (e.g., L-mannose) to natural “L-FimH” would, by symmetry, mirror the binding of D-FimH to D-mannose (or, conversely, the absence of such binding). Under this symmetry principle, binding of natural bacteria composed of L-proteins to L-glycans reflects the binding of mirror-image bacteria to natural D-sugars, thereby allowing evaluation of potential colonization risks in the human population.

*L. reuteri* and *E. coli*, two well-characterized gut-associated bacteria with established interactions with host glycans^36, 54^, were selected as model organisms to examine cross-chiral recognition of enantiomeric glycans by intact bacterial cells. We measured glycan binding of the *L. reuteri* JCM1081 with a LiGA containing enantiomeric monosaccharides-conjugated phages (**Figure 5A**), and a glycan display density of 4000 indicates that each phage carries at least one glycan per pVIII protein, with approximately half of the pVIII conjugated to two glycans. Surprisingly, *L. reuteri* JCM1081 exhibited a strong preference for L-glucose-[4000] over D-glucose (**Figure 5A**), despite L-glucose being rare in the poultry glycome. The JCM1081 strain also showed strong binding to both L- and D-fucose, whereas binding to L- and D-galactose-[4000] was comparatively weaker. One possible explanation for the enantiomer-independent recognition of fucose is the structural similarity between D-fucose (6-deoxy-D-galactose) and D-galactose and it has been demonstrated previously that *L. reuteri* JCM1081 exhibits strong binding to glycans terminating in D-Gal and L-Fuc such as Lac-peg4 and Di-N3 (**Figure 2A**).

**Figure 5.**
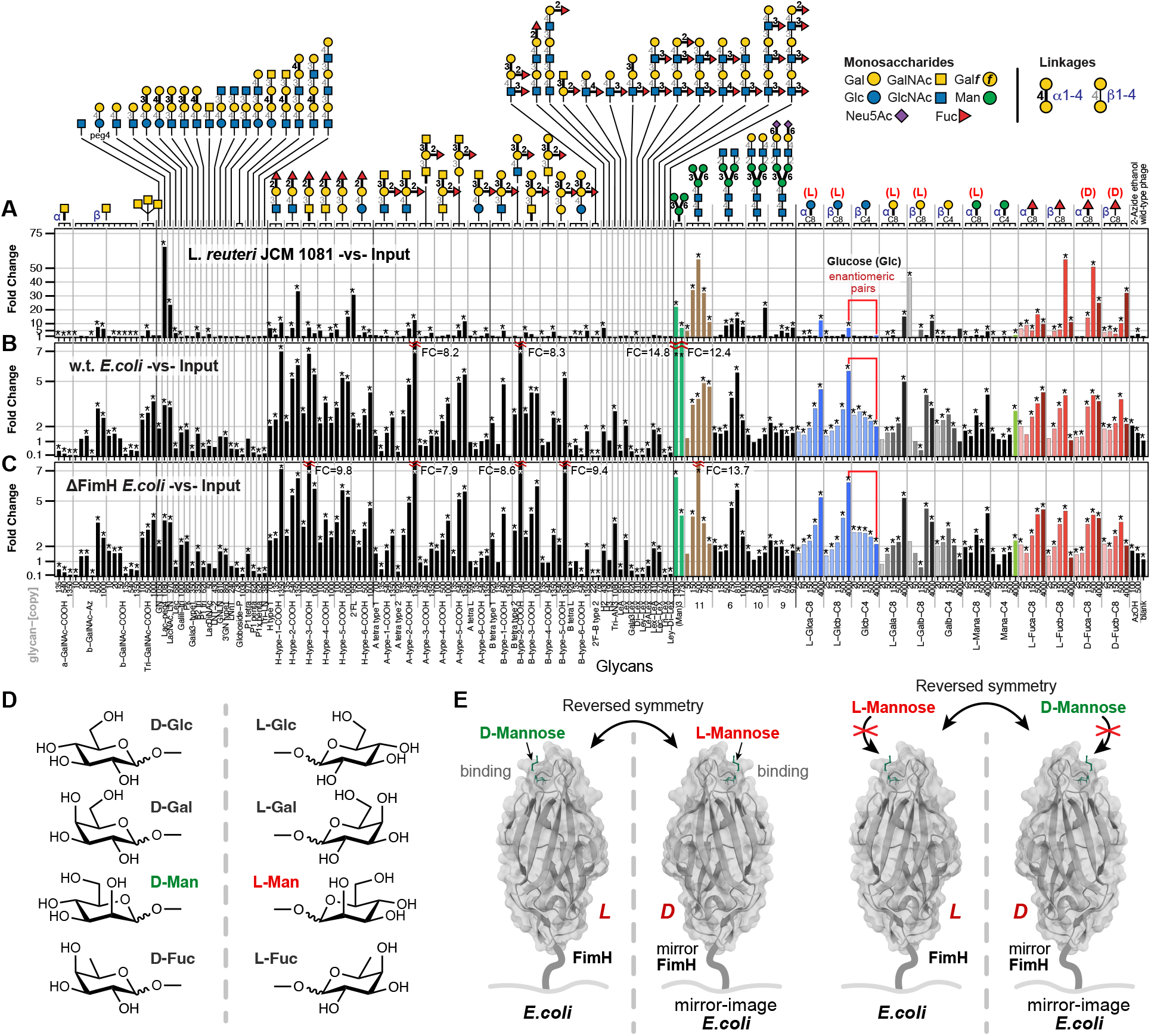
Evaluation of binding of natural monosaccharides and their mirror-image enantiomers to *L. reuteri* and *E. coli* with LiGA. (A) Glycan-binding profile of *L. reuteri* JCM1081 (n=6 for test samples). Display densities of enantiomeric pairs of glycans (L and D) were color-coded using shaded tones. (B) Glycan-binding profile of wild-type *E. coli* BW25113 (n = 5 for test samples). (C) Glycan-binding profile of FimH-deficient *E. coli* BW25113 (n = 5 for test samples). (D) Structure of natural monosaccharides (D-glucose, D-galactose, D-mannose, and L-fucose) and their enantiomers. (E) Natural proteins are composed exclusively of L-amino acids. FimH binds to D-Man, and by symmetry the mirror FimH binds to L-Man. By measuring binding of glycan enantiomers (e.g., L-Man) to natural “L-FimH” by symmetry measures the binding of D-FimH to D-Man (or in this case a lack of binding). By symmetry, binding of natural bacteria (made of L-proteins) to L-glycans also measures binding of mirror-image *E. coli* or mirror-image *L. reuteri* to natural D-sugars. Fold change, or FC, was calculated by Bioconductor edgeR DE analysis using the negative binomial model, TMM normalization and BH correction for FDR. All respective n values are independent biological replicates, and FC are reported with asterisks indicating FDR < 0.05. FimH (PDB 1TR7).

Glycan-binding profiling of the *E. coli* BW25113 with LiGA showed binding to α-D-Man-[Max] but not to the enantiomeric L-Man (**Figure 5B–C**). We also observed that both FimH^+^ and FimH^-^*E. coli* strain showed binding to L- and D-fucose in a density-dependent manner, with increasing the display density the binding increases, suggesting the cross-chiral recognition of enantiomeric pairs of Fuc is not FimH-dependent and the tested *E. coli* strains must express other molecules that mediated the binding. We also noted a minor but significant interaction between *E. coli* and L-glucose, and co-occurrence of L-fucose and L-glucose binding was in line with our recent observations that L-Glucose serves as effective mimetic of L-fucose^32^. In contrast, D-glucose, displayed at any density was not effective for binding to the *E. coli* tested.

## Discussion

Our findings revealed both conserved and strain-specific glycan-binding behaviors among bacteria, highlighting similarities within taxonomic groups and differences at the strain level. Pathogenic and commensal *E. coli* strains from humans and rodents displayed overlapping glycan-binding patterns, whereas *L. reuteri* and *Bacteroides* strains exhibited notable strain-dependent differences. Glycan-binding by *L. reuteri* has previously been characterized using the TLC overlay assay, inhibition assay by free carbohydrates and evaluating the different surface proteins of these bacteria. However, this paper represents the first comprehensive screening of live *L. reuteri* glycan-binding profiles using LiGA that contains diverse glycan structures with precisely controlled valency. Previous genetic analyses have shown that *L. reuteri* isolates cluster according to host species, and host specificity has been confirmed *in vivo*. Since glycan recognition often underlies host specificity in pathogens, we sought to evaluate whether glycan-binding of commensal *L. reuteri* strains is similarly host-dependent. Our results indicate that *L. reuteri* exhibits strain-specific, but not host-specific, glycan-binding behavior.

Across all 16 *L. reuteri* isolates tested, binding to mannose-containing glycans was consistently observed, which has been reported in some isolates. The *L. reuteri* 1063 (ATCC 53608) strain has been shown to bind mannose^55^, and the pig isolate *L. reuteri* lp167-67 bound exclusively to mannose-containing glycans, suggesting the presence of the mannose-specific adhesin (MSA) as its primary surface receptor. Mannose, an integral part of gut epithelial glycosylation, serves as a common adhesion target for microbes seeking epithelial colonization, and similar mechanism can be found in pathogens such as enterotoxigenic *E. coli* (ETEC) that express type I fimbriae to mediate mannose-specific adhesion to host cells^56^. Mannose-mediated adhesion has been implicated in the initial recognition of host tissues by *Campylobacter jejuni*^57^. Similarly, *Salmonella enterica* serovar Enteritidis, *Vibrio cholerae*, and *Pseudomonas aeruginosa* exploit mannose-containing glycoconjugates for host attachment^58^. Mannose-binding capability may contribute to the probiotic potential of *L. reuteri* by enabling competitive exclusion of pathogens through binding to mannose residues on the intestinal epithelium. The mannose binding capability of *L. reuteri* strains has been observed could mediate competitive inhibition of pathogenic bacteria within the intestine. It has been shown that adhesion of *Limosilactobacillus* to the epithelial layer via mannose-containing glycans can block *Salmonella* and ETEC attachment, thereby protecting the intestinal barrier^59^.

Surprisingly, we noticed two strains *L. reuteri* (rodent *L. reuteri* TMW1.656 and human *L. reuteri* DSM20016) exhibit binding to 3’SLN(Gc)-[570] that terminating with Neu5Gc. The glycocalyx of human cells differs from that of many other mammals, such as rodent, by the absence of Neu5Gc and an increased abundance of its precursor, Neu5Ac^60^. Uptake and incorporation of dietary Neu5Gc into salivary mucin O-glycans has been observed in humans^61^, and the adsorption of Neu5Gc from the diet may be the source of the O-glycans containing Neu5Gc.

Most *L. reuteri* (10/16) strains showed binding to the fucose-terminating glycans (Di-N3 and Le^x^) which suggests the presence of receptor(s) with fucose-binding activity and the presence of fucosylated glycans in their colonization site. It has been reported that the mucus-binding protein (MUB) mediated binding of *L. reuteri* ATCC53608 to mucus can be inhibited by the addition of fucose^62^ which the fucose-binding ability of the MUB. Similar inhibition assay can also be performed in future studies by competing LiGA binding to *L. reuteri* with excess free fucose or other glycans to confirm the presence of glycan-binding receptor(s).

The poultry-derived *L. reuteri* JCM1081 exhibited strong binding to Lac-peg4-[1080] and few other galactose containing glycans, consistent with a previous study reporting that the carbohydrate-binding activity of JCM1081 can be inhibited by galactose and lactose^63^. It has also been reported that *L. reuteri* JCM1081 can bind to asialo-GM1 (Galβ1-3GalNAcβ1-4Galβ1-4Glcβ1-1′Cer) and sulfatide (HSO3-3Galβ1-1′Cer) glycolipids^64^. The observation that *L. reuteri* JCM1081 showed strong binding to multiple glycans (e.g., Lac-peg4, LacNAc-LN, Lec, Di-N3, and 2’FL) in this study is also consistent with previous observations of its enhanced adhesive capacity to gut epithelial cells^34^. It has been reported that *L. reuteri* JCM1081 can inhibit binding of *Helicobacter pylori* to glycolipids receptors, including sulfatide, as demonstrated by thin-layer chromatography^64^, suggesting the importance of glycan binding of the *L. reuteri* in maintaining hemostasis and preventing the colonization of pathogenetic microbes. Overall, despite some similarities in glycan binding of *L. reuteri* strains, there is strain-specific glycan binding. Similar strain level glycan binding specificity has also been shown in the *Bacteroides* strains^17^.

The risk of mirror-imaged microorganisms has not been yet assessed in any evidence-based experiments because such organisms have not been produced yet. Glycans, with an incredibly prevalence in nature, can be found on the cell surface of all known life on earth. Here, we evaluated the colonization potential of mirror-imaged bacteria in host bearing natural chirality glycans by testing if natural isolated bacteria can engage with mirror-imaged glycans. We presented the very first evidence that the existence of cross-chiral recognition between naturally occurring bacteria and rare mirror-imaged glycans in the mammalian gut the microbes colonized. We found that D-fucose showed comparable binding to *E. coli* BW25113 as D-Man-C4 and *L. reuteri* JCM1081 exhibits a strong preference of L-glucose-[4000], which should be much rarely found in poultry gut, over the D-glucose. The JCM1081 strain also showed strong binding to L- and D-fucose and a relative weaker binding to L-/D-galactose-[4000]. Although only two bacterial strains has been tested in this study, we believe the cross-chiral recognition of glycans by microbes is more prevalent than expected.

Future work will be required to decipher the bacteria-glycan interactions in the temporal and spatial dynamics of GI tract mucus. Our study mapped glycan binding profiles of gut bacteria *in vitro* using LiGA, further *in vivo*—injection of LiGA in the gut and isolation of subtype of bacteria and associated LiGA—will identify the glycan-specific interactions and homing preferences in complex bacterial communities in the gut. Cross-chiral glycan binding capability of microbes will also be further investigated by mapping more bacterial strains using LiGA incorporated with a broader range of mirror-imaged complex glycans.

## Notes

The authors declare no competing interests. No unexpected or unusually high safety hazards were encountered in this study.

## Acknowledgments

The authors thank Prof. Jens Walter for kindly providing the *L. reuteri* strains and the staff at the University of Alberta mass spectrometry facility (Chemistry Department) for helping with MALDI analysis and S. Dang at the molecular biology service unit for assistance with Illumina sequencing.

## Supporting Information

Supplementary document contains Material and Methods, Supplementary Figures (S1-S6) and Supplementary Tables (S1-S5). Raw deep-sequencing data are publicly available at 48hd.cloud with data-specific URLs listed in **Table S5**.

## References

(1) Gao, J.; Liu, D.; Wang, Z. Screening lectin-binding specificity of bacterium by lectin microarray with gold nanoparticle probes. Anal. Chem. 2010, 82 (22), 9240–9247. Liu, Y.; Li, N.-q.; Zhao, X.-p.; Yue, B.; He, S.-w.; Gao, Z.-x.; Zhou, S.; Zhang, M. A C-type lectin that inhibits bacterial infection and facilitates viral invasion in black rockfish, Sebastes schlegelii. Fish Shellfish Immunol. 2016, 57, 309–317.

(2) Campanero-Rhodes, M. A.; Palma, A. S.; Menéndez, M.; Solís, D. Microarray strategies for exploring bacterial surface glycans and their interactions with glycan-binding proteins. Front. Microbiol. 2020, 10, 2909.

(3) Day, C. J.; Tran, E. N.; Semchenko, E. A.; Tram, G.; Hartley-Tassell, L. E.; Ng, P. S.; King, R. M.; Ulanovsky, R.; McAtamney, S.; Apicella, M. A. Glycan: glycan interactions: high affinity biomolecular interactions that can mediate binding of pathogenic bacteria to host cells. Proc. Natl. Acad. Sci. U.S.A. 2015, 112 (52), E7266–E7275.

(4) McPherson, R. L.; Isabella, C. R.; Walker, R. L.; Sergio, D.; Bae, S.; Gaca, T.; Raman, S.; Nguyen, L. T. T.; Wesener, D. A.; Halim, M. Lectin-Seq: a method to profile lectin-microbe interactions in native communities. Sci. Adv. 2023, 9 (30), eadd8766.

(5) Geissner, A.; Seeberger, P. H. Glycan arrays: from basic biochemical research to bioanalytical and biomedical applications. Annu. Rev. Anal. Chem. 2016, 9 (1), 223–247.

(6) Yan, M.; Zhu, Y.; Liu, X.; Lasanajak, Y.; Xiong, J.; Lu, J.; Lin, X.; Ashline, D.; Reinhold, V.; Smith, D. F. Next-generation glycan microarray enabled by DNA-coded glycan library and next-generation sequencing technology. Anal. Chem. 2019, 91 (14), 9221–9228.

(7) Lin, C.-L.; Sojitra, M.; Carpenter, E. J.; Hayhoe, E. S.; Sarkar, S.; Volker, E. A.; Wang, C.; Bui, D. T.; Yang, L.; Klassen, J. S. Chemoenzymatic synthesis of genetically-encoded multivalent liquid N-glycan arrays. Nat. Commun. 2023, 14 (1), 5237.

(8) Sojitra, M.; Sarkar, S.; Maghera, J.; Rodrigues, E.; Carpenter, E. J.; Seth, S.; Ferrer Vinals, D.; Bennett, N. J.; Reddy, R.; Khalil, A. Genetically encoded multivalent liquid glycan array displayed on M13 bacteriophage. Nat. Chem. Biol. 2021, 17 (7), 806–816.

(9) Sojitra, M.; Schmidt, E. N.; Lima, G. M.; Carpenter, E. J.; McCord, K. A.; Atrazhev, A.; Macauley, M. S.; Derda, R. Measuring carbohydrate recognition profile of lectins on live cells using liquid glycan array (LiGA). Nat. Protoc. 2025, 20 (4), 989–1019.

(10) Reddy, R.; Carpenter, E.; Halpin, A.; Sojitra, M.; Peng, C.; Lima, G. M.; Xue, X.; Yan, K.; Pearcy, J.; Ellis, M. Evaluation of Multiplexed Liquid Glycan Array (LiGA) for Serological Detection of Glycanbinding Antibodies. Glycobiology 2025, cwaf042.

(11) Dürr, C.; Nothaft, H.; Lizak, C.; Glockshuber, R.; Aebi, M. The Escherichia coli glycophage display system. Glycobiology 2010, 20 (11), 1366–1372. Celik, E.; Fisher, A. C.; Guarino, C.; Mansell, T. J.; DeLisa, M. P. A filamentous phage display system for N-linked glycoproteins. Protein Sci. 2010, 19 (10), 2006–2013. Çelik, E.; Ollis, A. A.; Lasanajak, Y.; Fisher, A. C.; Gür, G.; Smith, D. F.; DeLisa, M. P. Glycoarrays with engineered phages displaying structurally diverse oligosaccharides enable high-throughput detection of glycan–protein interactions. Biotechnol. J. 2015, 10 (1), 199–209.

(12) Pleiko, K.; Põšnograjeva, K.; Haugas, M.; Paiste, P.; Tobi, A.; Kurm, K.; Riekstina, U.; Teesalu, T. In vivo phage display: identification of organ-specific peptides using deep sequencing and differential profiling across tissues. Nucleic Acids Res. 2021, 49 (7), e38–e38.

(13) Arap, W.; Pasqualini, R. The human vascular mapping project. Selection and utilization of molecules for tumor endothelial targeting. Haemostasis 2001, 31, 30–31.

(14) Consortium, T. H. M. P. Structure, function and diversity of the healthy human microbiome. Nature 2012, 486 (7402), 207–214.

(15) Karcher, N.; Pasolli, E.; Asnicar, F.; Huang, K. D.; Tett, A.; Manara, S.; Armanini, F.; Bain, D.; Duncan, S. H.; Louis, P. Analysis of 1321 Eubacterium rectale genomes from metagenomes uncovers complex phylogeographic population structure and subspecies functional adaptations. Genome Biol. 2020, 21 (1), 138. Yaffe, E.; Relman, D. A. Tracking microbial evolution in the human gut using Hi-C reveals extensive horizontal gene transfer, persistence and adaptation. Nat. Microbiol. 2020, 5 (2), 343–353.

(16) Yang, C.; Mogno, I.; Contijoch, E. J.; Borgerding, J. N.; Aggarwala, V.; Li, Z.; Siu, S.; Grasset, E. K.; Helmus, D. S.; Dubinsky, M. C. Fecal IgA levels are determined by strain-level differences in Bacteroides ovatus and are modifiable by gut microbiota manipulation. Cell Host Microbe 2020, 27 (3), 467–475. e466.

(17) Patnode, M. L.; Guruge, J. L.; Castillo, J. J.; Couture, G. A.; Lombard, V.; Terrapon, N.; Henrissat, B.; Lebrilla, C. B.; Gordon, J. I. Strain-level functional variation in the human gut microbiota based on bacterial binding to artificial food particles. Cell Host Microbe 2021, 29 (4), 664–673. e665.

(18) Sun, S.; Tay, Q. X. M.; Kjelleberg, S.; Rice, S. A.; McDougald, D. Quorum sensing-regulated chitin metabolism provides grazing resistance to Vibrio cholerae biofilms. ISME J. 2015, 9 (8), 1812–1820.

(19) Miron, J.; Ben-Ghedalia, D.; Morrison, M. Invited review: adhesion mechanisms of rumen cellulolytic bacteria. J. Dairy Sci. 2001, 84 (6), 1294–1309.

(20) Poole, J.; Day, C. J.; von Itzstein, M.; Paton, J. C.; Jennings, M. P. Glycointeractions in bacterial pathogenesis. Nat. Rev. Microbiol. 2018, 16 (7), 440–452.

(21) Kalas, V.; Hibbing, M. E.; Maddirala, A. R.; Chugani, R.; Pinkner, J. S.; Mydock-McGrane, L. K.; Conover, M. S.; Janetka, J. W.; Hultgren, S. J. Structure-based discovery of glycomimetic FmlH ligands as inhibitors of bacterial adhesion during urinary tract infection. Proc. Natl. Acad. Sci. U.S.A. 2018, 115 (12), E2819–E2828. Le Guennec, L.; Virion, Z.; Bouzinba-Ségard, H.; Robbe-Masselot, C.; Leonard, R.; Nassif, X.; Bourdoulous, S.; Coureuil, M. Receptor recognition by meningococcal type IV pili relies on a specific complex N-glycan. Proc. Natl. Acad. Sci. U.S.A. 2020, 117 (5), 2606–2612.

(22) Dethlefsen, L.; McFall-Ngai, M.; Relman, D. A. An ecological and evolutionary perspective on human–microbe mutualism and disease. Nature 2007, 449 (7164), 811–818.

(23) Ley, R. E.; Hamady, M.; Lozupone, C.; Turnbaugh, P. J.; Ramey, R. R.; Bircher, J. S.; Schlegel, M. L.; Tucker, T. A.; Schrenzel, M. D.; Knight, R. Evolution of mammals and their gut microbes. Science 2008, 320 (5883), 1647–1651.

(24) Frese, S. A.; Benson, A. K.; Tannock, G. W.; Loach, D. M.; Kim, J.; Zhang, M.; Oh, P. L.; Heng, N. C.; Patil, P. B.; Juge, N. The evolution of host specialization in the vertebrate gut symbiont Lactobacillus reuteri. PLoS Genet. 2011, 7 (2), e1001314.

(25) Tannock, G. W. The lactic microflora of pigs, mice and rats. In The Lactic Acid Bacteria Volume 1: The Lactic Acid Bacteria in Health and Disease, Springer, 1992; pp 21–48. Leser, T. D.; Amenuvor, J. Z.; Jensen, T. K.; Lindecrona, R. H.; Boye, M.; Møller, K. Culture-independent analysis of gut bacteria: the pig gastrointestinal tract microbiota revisited. Appl. Environ. Microbiol. 2002, 68 (2), 673–690. Salzman, N. H.; de Jong, H.; Paterson, Y.; Harmsen, H. J.; Welling, G. W.; Bos, N. A. Analysis of 16S libraries of mouse gastrointestinal microflora reveals a large new group of mouse intestinal bacteria. Microbiology 2002, 148 (11), 3651–3660. Brooks, S. P.; McAllister, M.; Sandoz, M.; Kalmokoff, M. Culture-independent phylogenetic analysis of the faecal flora of the rat. Can. J. Microbiol. 2003, 49 (10), 589–601. Walter, J. Ecological role of lactobacilli in the gastrointestinal tract: implications for fundamental and biomedical research. Appl. Environ. Microbiol. 2008, 74 (16), 4985–4996.

(26) Jensen, H.; Roos, S.; Jonsson, H.; Rud, I.; Grimmer, S.; van Pijkeren, J.-P.; Britton, R. A.; Axelsson, L. Role of Lactobacillus reuteri cell and mucus-binding protein A (CmbA) in adhesion to intestinal epithelial cells and mucus in vitro. Microbiology 2014, 160 (4), 671–681.

(27) Kawai, Y.; Suegara, N. Specific adhesion of lactobacilli to keratinized epithelial cells of the rat stomach in vitro. Am. J. Clin. Nutr. 1977, 30 (11), 1777–1780. Schreiber, O.; Petersson, J.; Phillipson, M.; Perry, M.; Roos, S.; Holm, L. Lactobacillus reuteri prevents colitis by reducing P-selectin-associated leukocyte-and platelet-endothelial cell interactions. Am. J. Physiol. Gastrointest. Liver Physiol. 2009, 296 (3), G534–G542. Oh, P. L.; Benson, A. K.; Peterson, D. A.; Patil, P. B.; Moriyama, E. N.; Roos, S.; Walter, J. Diversification of the gut symbiont Lactobacillus reuteri as a result of host-driven evolution. ISME J. 2010, 4 (3), 377–387.

(28) Adamala, K. P.; Agashe, D.; Belkaid, Y.; Bittencourt, D. M. d. C.; Cai, Y.; Chang, M. W.; Chen, I. A.; Church, G. M.; Cooper, V. S.; Davis, M. M. Confronting risks of mirror life. Science 2024, 386 (6728), 1351–1353.

(29) Adamala, K.; Agashe, D.; Binder, D.; Cai, Y.; Cooper, V.; Duncombe, R.; Esvelt, K. Technical report on mirror bacteria: Feasibility and risks. Stanford Digital Repository: 2024.

(30) Derda, R. Remember The Glycans: Consideration of Glycans in Evaluating the Threat of Mirror-Image Life Forms. eLetter reply to “Confronting Risks of mirror life” Adamala et al. Science 2024, 386, 1351–1353.

(31) Varki, A. Biological roles of glycans. Glycobiology 2017, 27 (1), 3–49.

(32) Carpenter, E. J.; Peng, C.; Haregu, S.; Twells, N.; Woudstra, L.; Sood, A.; Cartmell, J.; Woods, R. J.; Mahal, L. K.; Wang, S.-K. Atom-level machine learning of protein-glycan interactions and cross-chiral recognition in glycobiology. Sci. Adv. 2025, 11 (49), eadx6373.

(33) Adler, J.; Hazelbauer, G. L.; Dahl, M. Chemotaxis toward sugars in Escherichia coli. J. Bacteriol. 1973, 115 (3), 824–847.

(34) Wang, B.; Wei, H.; Yuan, J.; Li, Q.; Li, Y.; Li, N.; Li, J. Identification of a surface protein from Lactobacillus reuteri JCM1081 that adheres to porcine gastric mucin and human enterocyte-like HT-29 cells. Curr. Microbiol. 2008, 57 (1), 33–38.

(35) Brockhausen, I.; Dowler, T.; Paulsen, H. Site directed processing: role of amino acid sequences and glycosylation of acceptor glycopeptides in the assembly of extended mucin type O-glycan core 2. Biochim. Biophys. Acta Gen. Subj. 2009, 1790 (10), 1244–1257. Karlsson, N. G.; Nordman, H.; Karlsson, H.; Carlstedt, I.; Hansson, G. C. Glycosylation differences between pig gastric mucin populations: a comparative study of the neutral oligosaccharides using mass spectrometry. Biochem. J. 1997, 326 (3), 911–917. Gunning, A. P.; Kirby, A. R.; Fuell, C.; Pin, C.; Tailford, L. E.; Juge, N. Mining the “glycocode”—exploring the spatial distribution of glycans in gastrointestinal mucin using force spectroscopy. FASEB J. 2013, 27 (6), 2342.

(36) Tassell, M. L. V.; Miller, M. J. Lactobacillus adhesion to mucus. Nutrients 2011, 3 (5), 613–636.

(37) Park, D.; Xu, G.; Barboza, M.; Shah, I. M.; Wong, M.; Raybould, H.; Mills, D. A.; Lebrilla, C. B. Enterocyte glycosylation is responsive to changes in extracellular conditions: implications for membrane functions. Glycobiology 2017, 27 (9), 847–860. Gebhard, A.; Gebert, A. Brush cells of the mouse intestine possess a specialized glycocalyx as revealed by quantitative lectin histochemistry: further evidence for a sensory function. J. Histochem. Cytochem. 1999, 47 (6), 799–808.

(38) Alverdy, J.; Bergelson, J.; Liu, D.; Poroyko, V.; Morowitz, M.; Bell, T.; Ulanov, A.; Wang, M.; Donovan, S.; Bao, N. Diet creates metabolic niches in the” inmature gut” that shape microbial communities. Nutr. Hosp. 2011, 26 (6), 1283–1295.

(39) Martens, E. C.; Chiang, H. C.; Gordon, J. I. Mucosal glycan foraging enhances fitness and transmission of a saccharolytic human gut bacterial symbiont. Cell Host Microbe 2008, 4 (5), 447–457.

(40) Phansopa, C.; Roy, S.; Rafferty, J. B.; Douglas, C. I.; Pandhal, J.; Wright, P. C.; Kelly, D. J.; Stafford, G. P. Structural and functional characterization of NanU, a novel high-affinity sialic acid-inducible binding protein of oral and gut-dwelling Bacteroidetes species. Biochem. J. 2014, 458 (3), 499–511.

(41) Coker, J. K.; Moyne, O.; Rodionov, D. A.; Zengler, K. Carbohydrates great and small, from dietary fiber to sialic acids: How glycans influence the gut microbiome and affect human health. Gut Microbes 2021, 13 (1), 1869502.

(42) Berne, C.; Ellison, C. K.; Ducret, A.; Brun, Y. V. Bacterial adhesion at the single-cell level. Nat. Rev. Microbiol. 2018, 16 (10), 616–627. Xu, Q.; Shoji, M.; Shibata, S.; Naito, M.; Sato, K.; Elsliger, M.-A.; Grant, J. C.; Axelrod, H. L.; Chiu, H.-J.; Farr, C. L. A distinct type of pilus from the human microbiome. Cell 2016, 165 (3), 690–703.

(43) Rossez, Y.; Gosset, P.; Boneca, I. G.; Magalhães, A.; Ecobichon, C.; Reis, C. A.; Cieniewski-Bernard, C.; Joncquel Chevalier Curt, M.; Léonard, R.; Maes, E. The lacdiNAc-specific adhesin LabA mediates adhesion of Helicobacter pylori to human gastric mucosa. J. Infect. Dis 2014, 210 (8), 1286–1295.

(44) Wyatt, J.; Rathbone, B.; Sobala, G.; Shallcross, T.; Heatley, R.; Axon, A.; Dixon, M. Gastric epithelium in the duodenum: its association with Helicobacter pylori and inflammation. J. Clin. Pathol. 1990, 43 (12), 981–986.

(45) Van de Vijver, K. K.; Deelder, A. M.; Jacobs, W.; Van Marck, E. A.; Hokke, C. H. LacdiNAc- and LacNAc-containing glycans induce granulomas in an in vivo model for schistosome egg-induced hepatic granuloma formation. Glycobiology 2006, 16 (3), 237–243.

(46) Haga, Y.; Uemura, M.; Baba, S.; Inamura, K.; Takeuchi, K.; Nonomura, N.; Ueda, K. Identification of multisialylated LacdiNAc structures as highly prostate cancer specific glycan signatures on PSA. Anal. Chem. 2019, 91 (3), 2247–2254.

(47) Yutin, N.; Galperin, M. Y. A genomic update on clostridial phylogeny: G ram-negative spore formers and other misplaced clostridia. Environ. Microbiol 2013, 15 (10), 2631–2641.

(48) Gelberg, H. B. Alimentary system and the peritoneum, omentum, mesentery, and peritoneal cavity. Pathologic basis of veterinary disease 2017, 324.

(49) Bouckaert, J.; Mackenzie, J.; De Paz, J. L.; Chipwaza, B.; Choudhury, D.; Zavialov, A.; Mannerstedt, K.; Anderson, J.; Piérard, D.; Wyns, L. The affinity of the FimH fimbrial adhesin is receptor-driven and quasi-independent of Escherichia coli pathotypes. Mol. Microbiol. 2006, 61 (6), 1556–1568.

(50) Day, C. J.; Lo, A. W.; Hartley-Tassell, L. E.; Argente, M. P.; Poole, J.; King, N. P.; Tiralongo, J.; Jennings, M. P.; Schembri, M. A. Discovery of bacterial fimbria–glycan interactions using whole-cell recombinant Escherichia coli expression. mBio 2021, 12 (1), 10.1128/mbio.03664-03620.

(51) Mottram, L.; Liu, J.; Chavan, S.; Tobias, J.; Svennerholm, A.-M.; Holgersson, J. Glyco-engineered cell line and computational docking studies reveals enterotoxigenic Escherichia coli CFA/I fimbriae bind to Lewis a glycans. Sci. Rep. 2018, 8 (1), 11250.

(52) Coddens, A.; Diswall, M.; Ångström, J.; Breimer, M. E.; Goddeeris, B.; Cox, E.; Teneberg, S. Recognition of blood group ABH type 1 determinants by the FedF adhesin of F18-fimbriated Escherichia coli. J. Biol. Chem. 2009, 284 (15), 9713–9726.

(53) Esko, J. D.; Sharon, N. Microbial Lectins: Hemagglutinins, Adhesins, and Toxins. In Essentials of Glycobiology, Varki, A., Cummings, R. D., Esko, J. D., Freeze, H. H., Stanley, P., Bertozzi, C. R., Hart, G. W., Etzler, M. E. Eds.; Cold Spring Harbor Laboratory Press Copyright © 2009, The Consortium of Glycobiology Editors, La Jolla, California., 2009.

(54) Luis, A. S.; Hansson, G. C. Intestinal mucus and their glycans: A habitat for thriving microbiota. Cell Host Microbe 2023, 31 (7), 1087–1100.

(55) Roos, S.; Jonsson, H. A high-molecular-mass cell-surface protein from Lactobacillus reuteri 1063 adheres to mucus components. Microbiology 2002, 148 (2), 433–442.

(56) Reid, G.; Burton, J. Use of Lactobacillus to prevent infection by pathogenic bacteria. Microbes Infect. 2002, 4 (3), 319–324.

(57) Day, C. J.; Tram, G.; Hartley-Tassell, L. E.; Tiralongo, J.; Korolik, V. Assessment of glycan interactions of clinical and avian isolates of Campylobacter jejuni. BMC Microbiol. 2013, 13 (1), 228.

(58) Bhattacharjee, J.; Srivastava, B. S. Adherence of wild-type and mutant strains of Vibrio cholerae to normal and immune intestinal tissue. Bull. World Health Organ. 1979, 57 (1), 123. Imberty, A.; Wimmerová, M.; Mitchell, E. P.; Gilboa-Garber, N. Structures of the lectins from Pseudomonas aeruginosa: insights into the molecular basis for host glycan recognition. Microbes Infect. 2004, 6 (2), 221–228.

(59) Yue, M.; Rankin, S. C.; Blanchet, R. T.; Nulton, J. D.; Edwards, R. A.; Schifferli, D. M. Diversification of the Salmonella fimbriae: a model of macro-and microevolution. PLoS One 2012, 7 (6), e38596. Garcia-Gonzalez, N.; Prete, R.; Battista, N.; Corsetti, A. Adhesion properties of food-associated Lactobacillus plantarum strains on human intestinal epithelial cells and modulation of IL-8 release. Front. Microbiol. 2018, 9, 2392.

(60) Altman, M. O.; Gagneux, P. Absence of Neu5Gc and presence of anti-Neu5Gc antibodies in humans—an evolutionary perspective. Front. Immunol. 2019, 10, 789.

(61) Tangvoranuntakul, P.; Gagneux, P.; Diaz, S.; Bardor, M.; Varki, N.; Varki, A.; Muchmore, E. Human uptake and incorporation of an immunogenic nonhuman dietary sialic acid. Proc. Natl. Acad. Sci. U.S.A. 2003, 100 (21), 12045–12050.

(62) MacKenzie, D. A.; Jeffers, F.; Parker, M. L.; Vibert-Vallet, A.; Bongaerts, R. J.; Roos, S.; Walter, J.; Juge, N. Strain-specific diversity of mucus-binding proteins in the adhesion and aggregation properties of Lactobacillus reuteri. Microbiology 2010, 156 (11), 3368–3378.

(63) Mukai, T.; Kaneko, S.; Ohori, H. Haemagglutination and glycolipid-binding activities of Lactobacillus reuteri. Lett. Appl. Microbiol. 1998, 27 (3), 130–134.

(64) Mukai, T.; Asasaka, T.; Sato, E.; Mori, K.; Matsumoto, M.; Ohori, H. Inhibition of binding of Helicobacter pylori to the glycolipid receptors by probiotic Lactobacillus reuteri. FEMS Immunol. Med. Microbiol. 2002, 32 (2), 105–110.

