## Supplementary figures and tables for "Mapping Glycan Binding Profiles of the Gut Microbes using Liquid Glycan Array (LiGA)"

### 1. Methods

#### a). Bacterial culturing

To determine whether glycan-binding profiles were similar among *L. reuteri* strains from the same host, we analyzed glycan binding in 16 *L. reuteri* strains, comprising four isolates each from murine, porcine, poultry, and human lineages (**Table S2**). All bacteria were grown in the media indicated in **Table S3**. *L. reuteri* strains were cultured in de Man, Rogosa, and Sharpe (MRS) medium (Difco) supplemented with 5% maltose and 10% fructose under anaerobic conditions (5% $CO_2$, 5% $H_2$, and 90% $N_2$) in anaerobic chamber. All *E. coli* strains were grown in Luria-Bertani (LB) broth with agitation. Each strain was cultured, and optical density at 600 nm (OD600) was measured using NanoDrop (Thermo Fisher Scientific), with CFU $mL^{-1}$ determined at 12, 18, and 24 h (**Table S4**).

#### b). Sanger sequencing

Each bacterial strain was tested by the colony PCR of 16S rRNA gene regions using 8F and 926R primers, followed by Sanger sequencing[1]. PCR reactions (50 µl) contained 2 µl of each primer (10 µM), 2 µl dNTP mix (10 mM; Invitrogen), 5 µl 10× Taq polymerase buffer (Invitrogen), 2 µl $MgCl_2$ (50 mM; Invitrogen), 0.5 µl Taq polymerase (1 U $\mu l^{-1}$; Invitrogen), and a small amount of bacterial colony. Thermal cycling consisted of an initial denaturation at 94 °C for 10 min, followed by 40 cycles of 94 °C for 30 s, 56 °C for 30 s, and 72 °C for 1 min, with a final extension at 72 °C for 7 min. PCR products were resolved by 1% agarose gel electrophoresis and visualized using SYBR Safe DNA gel stain (Invitrogen). Sequence identities were confirmed using nucleotide BLAST.

#### c). Quantifying phage titer by plaque-forming assay or quantitative PCR (qPCR)

Titer of phage in the input LiGA library and phages bound to bacteria in LiGA binding assays were quantified by plaque-forming assay. Briefly, the host *E. coli* K-12 strain was cultured overnight in Luria-Bertani (LB) medium with shaking at 37 °C. For phage titering, samples containing phage were serially diluted in HEPES buffer (50 mM HEPES, 150 mM NaCl, 2 mM $CaCl_2$, 10 mM $MgCl_2$), and 10 µl of each dilution was mixed with 200 µl of *E. coli* K-12 culture and 3 ml of molten top agar. The mixture was poured onto LB agar plates supplemented with X-gal and IPTG (40 µg $ml^{-1}$ each). After solidification, plates were incubated overnight at 37 °C, and phage titers (PFU $ml^{-1}$) were calculated by counting plaques formed on agar plates.

Alternatively, phage titer was determined by qPCR. Briefly, phage sample (2 µl) was mixed with universal qPCR mix (10 µl, product of the MBSU at University of Alberta), qPCR primer mix (5.5 uL, 2 µM each), and nuclease-free water (2.5 µl, IDT), followed by quantification using the CFX96 Touch Real-Time PCR Detection System (Bio-Rad). The phage titer was converted from Cq value using a calibration curve which determined by qPCR quantifying serial diluted phage sample with a known initial titer.

#### d). LiGA preparation

LiGA was prepared by following a previously reported procedure[2]. Glycan structures present in the LiGA-ED are listed in **Table S1**. Generation and characterization by MALDI-TOF MS of individual LiGA components can be found in our previous publications[2, 3, 4].

**e). Profiling glycan binding of bacterial cells using LiGA**

The LiGA-binding assay was performed following a protocol adapted from our previous publications[2, 4]. For each bacterial strain, biological replicates were prepared by inoculating single colony in 5 ml of broth media (e.g., five colonies were cultured individually for five downstream LiGA-binding assays). After culturing, $OD_{600}$ was quantified using a Nanodrop spectrophotometer (Thermo Fisher Scientific), and a volume of culture containing $1\times10^8$ CFU of bacteria were pelleted (5 min, 9,391 ×g). Bacteria pellet was washed once with 1 ml HEPES buffer to remove residual medium.

For LiGA binding, washed bacterial pellets were gently resuspended in 100 µl HEPES binding buffer containing LiGA at a final concentration of $1\times10^9$ PFU $ml^{-1}$. Samples were incubated on ice for 1 h with gentle mixing every 15 min. Following incubation, bacteria were pelleted by centrifugation (2 min, 9,391 × g) and washed three times with 1 ml ice-cold HEPES binding buffer. After the final wash, phage DNA was extracted and purified using the GeneJET Plasmid Miniprep Kit (Thermo Fisher Scientific). Purified DNA was used directly for qPCR and indexing PCR.

**f). PCR amplification of the bound glycophages to bacteria from LiGA**

The extracted DNA was subjected to a two-step PCR amplification procedure immediately after LiGA binding assay.

In the first-step PCR, the DNA template (20 µl) was amplified in a total reaction volume of 50 µl with 5x HF buffer (10 µl, NEB, Catalog# M0530S), 10 mM dNTP mix (1 µl, ThermoFisher Scientific), PCR primer mix (5 µl, 10 µM each), DMSO (1 µl), Phusion™ High-Fidelity DNA Polymerase (0.5 µl, NEB, Catalog# M0530S), and nuclease-free water (12.5 µl). In amplification of naïve LiGA, volume of DNA template was 2 µl. Thermal cycling was performed using the following settings: a) 95 °C for 5 min, b) 95 °C for 10 s, c) 50 °C for 30 s, d) 72 °C for 20 s, e) repeat b-d for 30 cycles, f) 4 °C hold.

The second-step PCR adds Illumina indexing barcodes to the DNA product of the first-step PCR. The DNA template (5 µl) was amplified in a total reaction volume of 50 µl with 5x HF buffer (10 µl, NEB, Catalog# M0530S), 10 mM dNTP mix (1 µl, ThermoFisher Scientific), forward SDB indexing primer (2.5 µl, 10 µM), reverse SDB indexing primer (2.5 µl, 10 µM), Phusion™ High-Fidelity DNA Polymerase (0.5 µl, NEB, Catalog# M0530S), and nuclease-free water (28.5 µl). Thermal cycling was performed using the following settings: a) 95 °C for 5 min, b) 95 °C for 10 s, c) 50 °C for 30 s, d) 72 °C for 20 s, e) repeat b-d for 15 cycles, f) 4 °C hold. PCR products were resolved by 2% agarose gel electrophoresis to confirm complete of PCR amplification, and indexed-DNA samples were submitted for deep sequencing[2]. Processing of the data is described below.

**g). Processing of Illumina data**

The procedure of Illumina data processing was adapted from our previous publication[2]. The Gzip compressed FASTQ files were downloaded from BaseSpace™ Sequence Hub. The files were converted into tables of DNA sequences and their counts per experiment. Briefly, FASTQ files were parsed into separate files based on unique multiplexing barcodes within the reads. Reads that did not contain an identifiable multiplex barcode were discarded. Several quality control steps were performed based on i) reads that contained a Phred quality score = 0 in any position were also discarded (ii) mapping the forward and reverse primer regions was done allowing no more than one base substitution each, (iii) alignment of the forward and reverse read-end overlap was performed allowing no mismatches between forward and reverse read in the overlap region. Reads that did not match criteria (ii) and (iii) were discarded. The two ends of read-pairs that pass the filtering criteria were joined and trimmed to the DNA sequences located between the priming regions; the reads were organized in a tab-delimited text file containing the unique DNA sequences and their copy numbers. Technical replicates were combined in the same file. Using an SDB-lookup table, DNA sequences were mapped to SDB and were translated to glycans using a LiGA-specific lookup table ("LiGA dictionary"). The file with DNA reads, raw counts, and mapped glycans, were uploaded to https://48hd.cloud/ server. Each experiment has a unique alphanumeric name (e.g., 20210813-87EDcsfGT-OB). All LiGA binding data used in this study has been listed in **Table S5**.

**h). Data analysis**

Data analysis was performed in R–Bioconductor and was adapted from our previous publication[2,4]. Comparisons and testing differences for significance in the LiGA data were performed as described in publication[5] using DE analysis implemented in edgeR[6]. During the DE analysis, three factors were considered: (1) modeling of the observed counts using a negative binomial model, (2) BH adjustment to control the FDR at $\alpha = 0.05$[7], and (3) normalization of data across multiple samples using TMM normalization[8]. To assess the significance of glycan binding in a specific experiment, the DE of the levels of the DNA barcode associated with that glycan in 'test' sets of the DNA read was compared to that of the levels of the same read in 'control' sets. For example, in bacterial cells, the 'test' dataset was association of the LiGA with the cells, whereas the control was naïve library. For binding to lectins, the 'control' dataset was association of the LiGA with blank carriers (blank wells in plates). Before DE analysis, 'test' and 'control' datasets were retrieved from the server at https://48hd.cloud/ as tables of glycans, DNA and raw sequencing counts (**Table S5**).

### 2. Figures and Tables Referenced in the Main Text

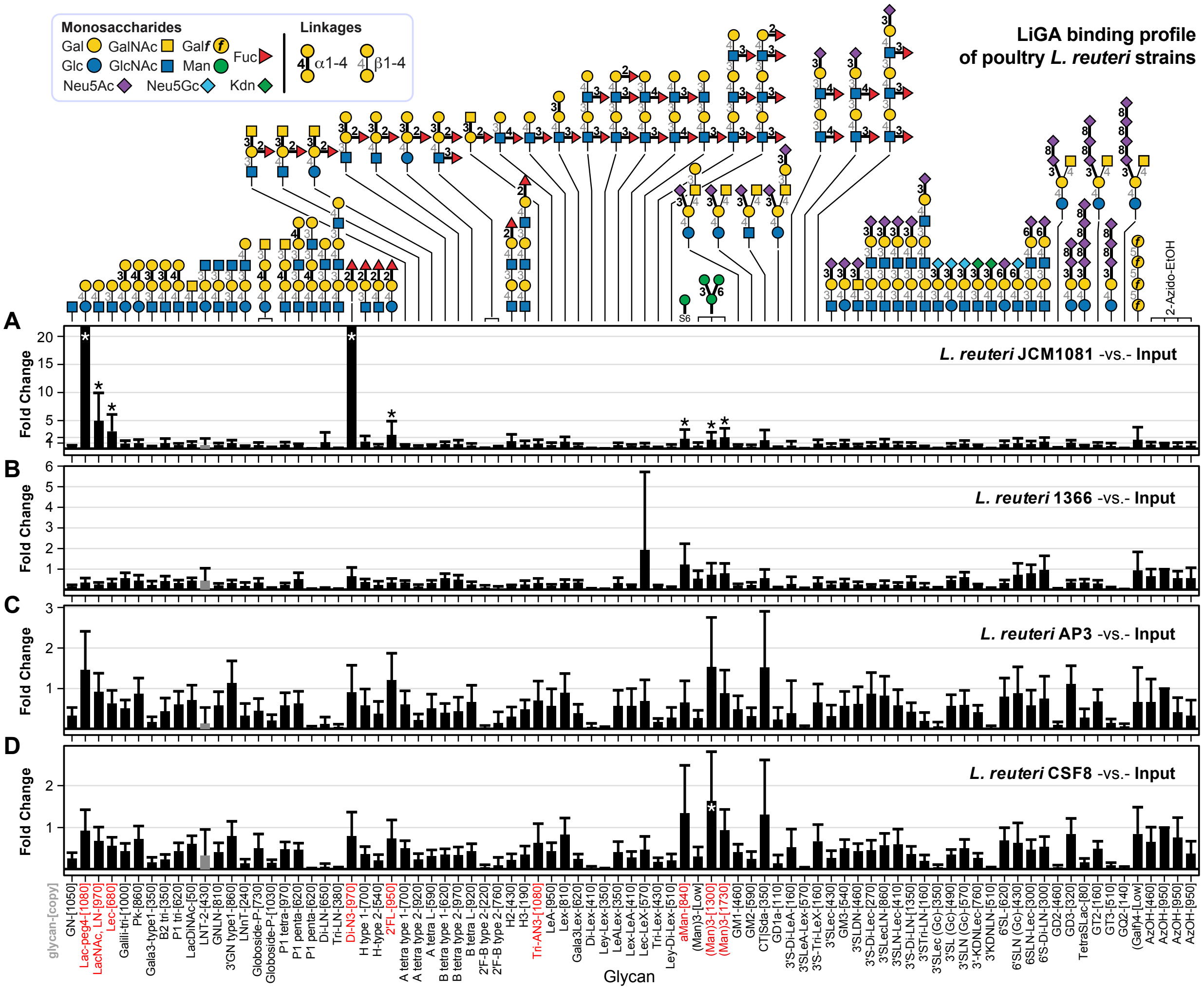

**Figure S1. Glycan binding of the poultry *L. reuteri* strains.**
Fold change (FC) was calculated by Bioconductor edgeR DE analysis using the negative binomial model, TMM normalization and BH correction for False-Discovery Rate (FDR). Error bars represent s.d. propagated from the variance of the TMM-normalized sequencing data. Glycan notations and color codes of α and β linkages are shown in the legend. * represents FDR ≤ 0.05, n = 7.

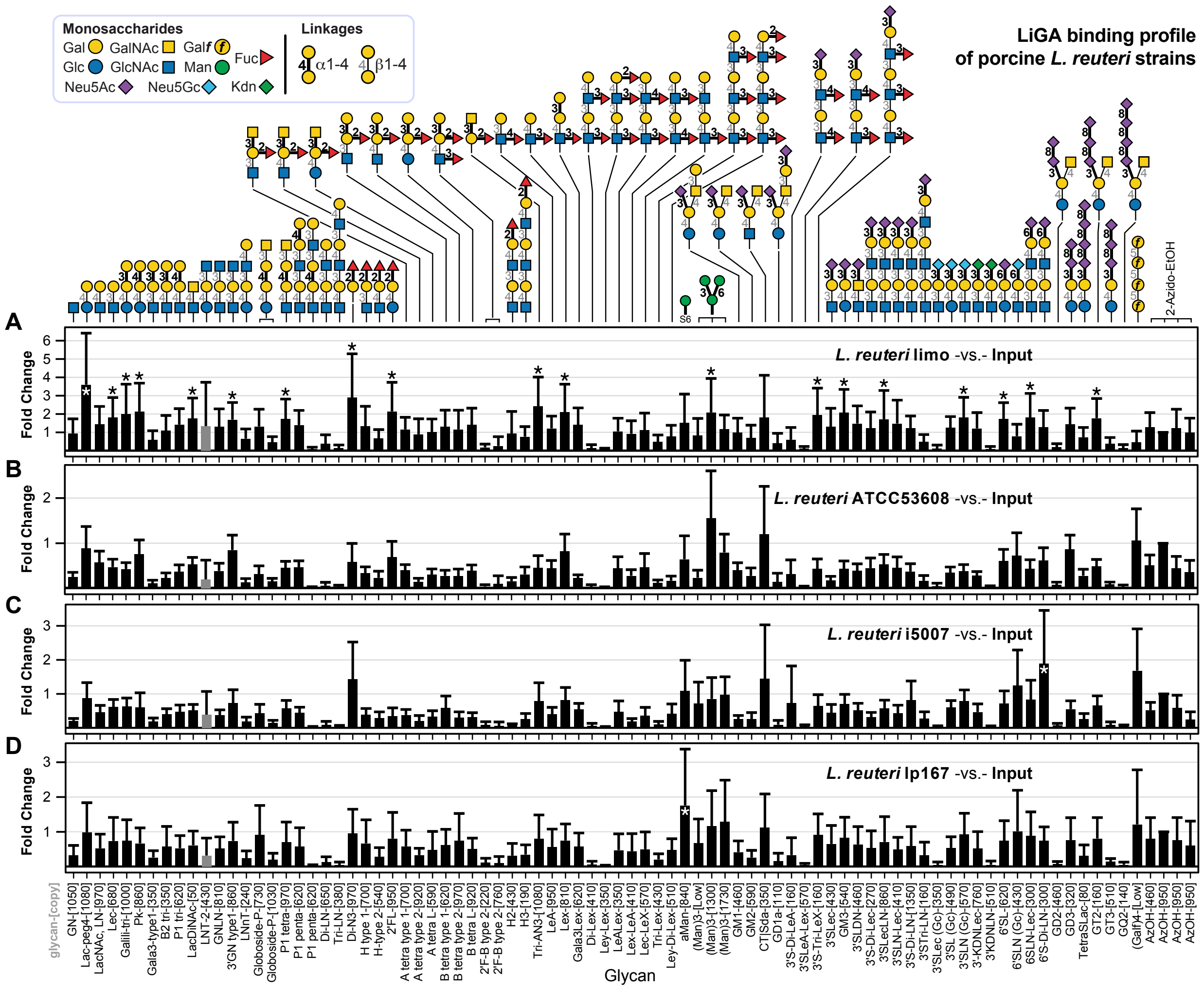

**Figure S2. Glycan binding of the porcine *L. reuteri* strains.**

FC was calculated by Bioconductor edgeR DE analysis using the negative binomial model, TMM normalization and BH correction for FDR. Error bars represent s.d. propagated from the variance of the TMM-normalized sequencing data. Glycan notations and color codes of α and β linkages are shown in the legend. * represents FDR ≤ 0.05, n = 7.

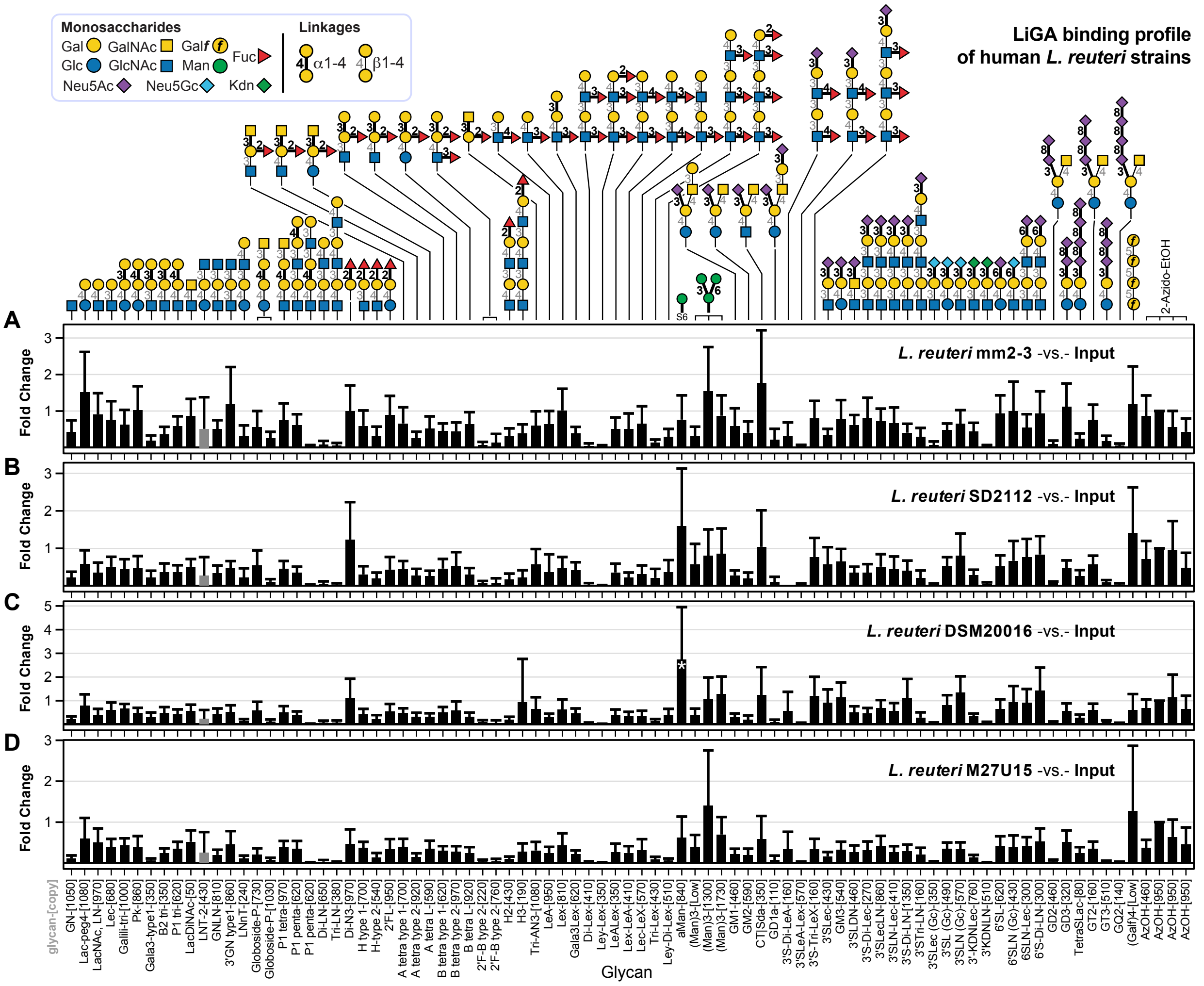

**Figure S3. Glycan binding of the human *L. reuteri* strains.**

FC was calculated by Bioconductor edgeR DE analysis using the negative binomial model, TMM normalization and BH correction for FDR. Error bars represent s.d. propagated from the variance of the TMM-normalized sequencing data. Glycan notations and color codes of α and β linkages are shown in the legend. * represents FDR ≤ 0.05, n = 7.

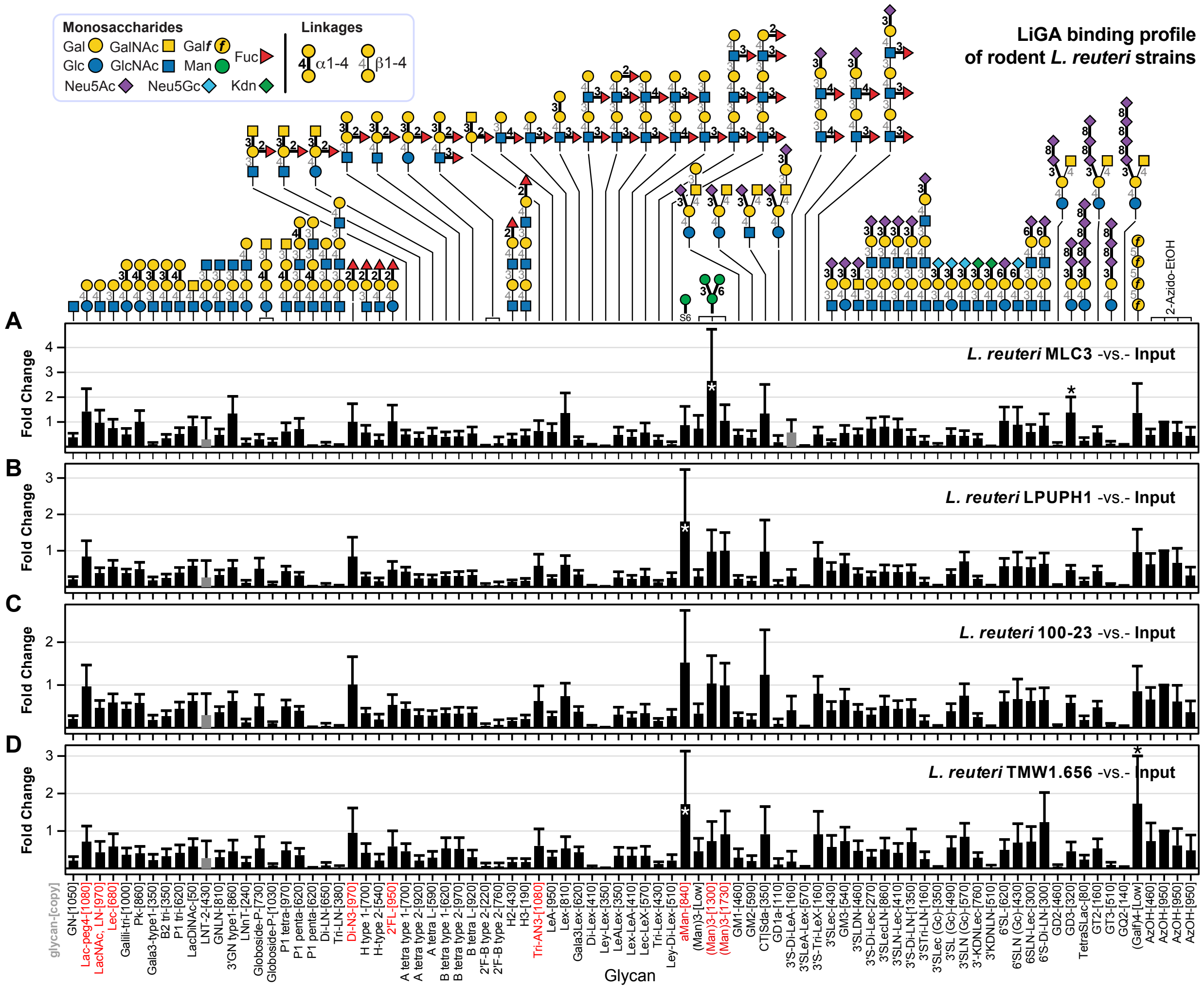

**Figure S4. Glycan binding of the rodent *L. reuteri* strains.**
FC was calculated by Bioconductor edgeR DE analysis using the negative binomial model, TMM normalization and BH correction for FDR. Error bars represent s.d. propagated from the variance of the TMM-normalized sequencing data. Glycan notations and color codes of α and β linkages are shown in the legend. * represents FDR ≤ 0.05, n = 7.

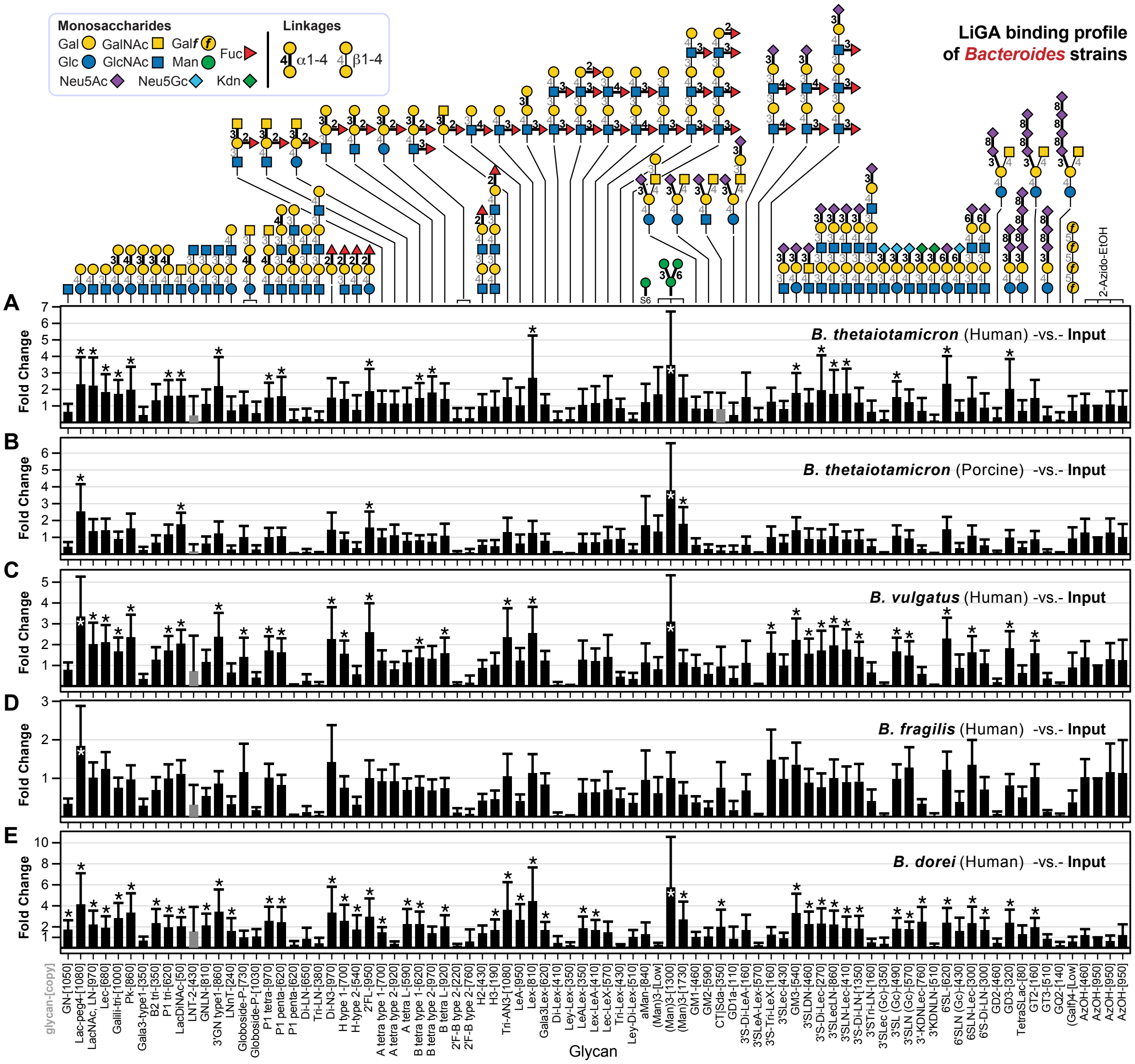

**Figure S5. Glycan binding of the *Bacteroides* strains.**

FC was calculated by Bioconductor edgeR DE analysis using the negative binomial model, TMM normalization and BH correction for FDR. Error bars represent s.d. propagated from the variance of the TMM-normalized sequencing data. * represents FDR ≤ 0.05, n = 7. a) *Bacteroides theiotamicron* human strain, b) *Bacteroides theiotamicron* pig strain c) *Bacteroides vulgatus* human strain, d) *Bacteroides fragilis* and e) *Bacteroides dorei* human strain.

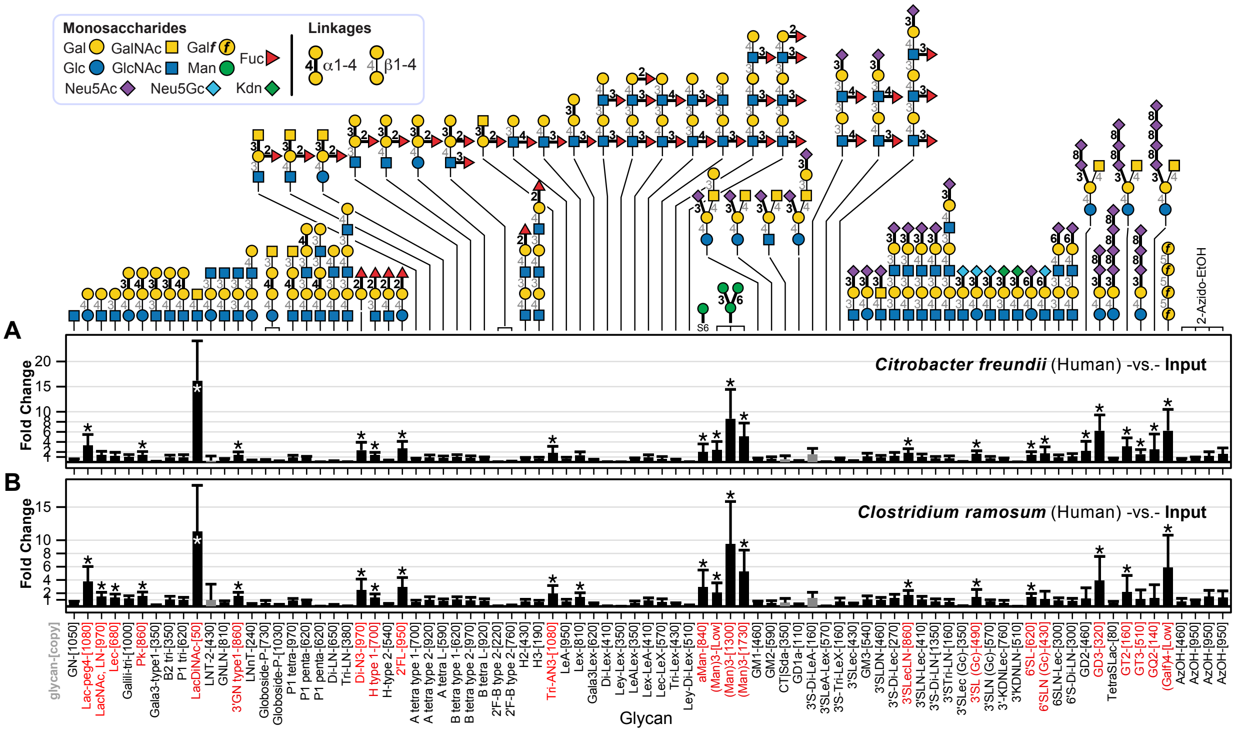

**Figure S6. Glycan binding of *C. freudii* and *C. ramosum*.**
FC was calculated by Bioconductor edgeR DE analysis using the negative binomial model, TMM normalization and BH correction for FDR. Error bars represent s.d. propagated from the variance of the TMM-normalized sequencing data. * represents FDR ≤ 0.05, n = 7. a) *Citrobacter freundii* human strain, b) *Clostridium ramosum* (*Erysipelatoclostridium ramosum*) human strain.

| Common name | Display density (glycan per phage) | Glycan chemical structure | LiGA-ED | LiGA-NOA |
|---|---|---|---|---|
| a-GalNAc-COOH | [135], [540], [1350] | GalNAcα-O-butyl-COOH | | |
| b-GalNAc-Az | [10], [20], [50], [100], [500], [1000] | GalNAc(β-P3 | | |
| b-GalNAc-COOH | [10], [20], [50], [100], [500], [1000] | GalNAcβ-O-PEG4-COOH | | |
| Tri-GalNAc-COOH | [100], [500], [1000] | (β-D-GalNAc-sp)3-NHCO-PEG5-COOH | | |
| GN | [1050] | GlcNAc(β-Sp | | |
| Lac-peg4 | [1080] | Galβ1-4Glc(β-P4 | | |
| LacNAc, LN | [970] | Galβ1-4GlcNAc(β-Sp | | |
| Lec | [680] | Galβ1-3GlcNAc(β-Sp | | |
| Galili-tri | [1000] | Galα1-3Galβ1-4Glc(β-Sp | | |
| Pk | [860] | Galα1-4Galβ1-4Glc(β-Sp | | |
| Gala3-type1 | [350] | Galα1-3Galβ1-3GlcNAc(β-Sp | | |
| B2 tri | [350] | Galα1-3Galβ1-4GlcNAc(β-Sp | | |
| P1 tri | [620] | Galα1-4Galβ1-4GlcNAc(β-Sp | | |
| LacDiNAc | [50] | GalNAcβ1-4GlcNAc(β-Sp | | |
| LNT-2 | [430] | GlcNAcβ1-3Galβ1-4Glc(β-Sp | | |
| GNLN | [810] | GlcNAcβ1-3Galβ1-4GlcNAc(β-Sp | | |
| 3'GN type1 | [860] | GlcNAcβ1-3Galβ1-3GlcNAc(β-Sp | | |
| LNnT | [240] | Galβ1-4GlcNAcβ1-3Galβ1-4Glc(β-Sp | | |
| Globoside-P | [730], [1030] | GalNAcβ1-3Galα1-4Galβ1-4Glc(β-Sp | | |
| P1 tetra | [970] | GalNAcβ1-3Galα1-4Galβ1-4GlcNAc(β-Sp | | |
| P1 penta | [620] | Galα1-4Galβ1-4GlcNAcβ1-3Galβ1-4Glc(β-Sp | | |
| P1x penta | [620] | Galβ1-3GalNAcβ1-3Galα1-4Galβ1-4GlcNAc(β-Sp | | |
| Di-LN | [650] | Galβ1-4GlcNAcβ1-3Galβ1-4GlcNAcβ1-3(β-Sp | | |

| | | |
|---|---|---|
| Tri-LN | [380] | Galβ1-4GlcNAcβ1-3Galβ1-4GlcNAcβ1-3Galβ1-4GlcNAcβ1-3(β-Sp |
| Di-N3 | [970] | Fucα1-2Gal(β-Sp |
| H type 1 | [700] | Fucα1-2Galβ1-3GlcNAc(β-Sp |
| H-type-1-COOH | [135], [1350] | Fucα1-2Galβ1-3GlcNAc(β-COOH |
| H-type-2-[540] | [540] | Fucα1-2Galβ1-4GlcNAc(β-Sp |
| H-type-2-COOH | [135], [540], [1350] | Fucα1-2Galβ1-4GlcNAc(β-COOH |
| H-type-3-COOH | [100], [500], [1000] | Fucα1-2Galβ1-3GalNAc(α-COOH |
| H-type-4-COOH | [100], [500], [1000] | Fucα1-2Galβ1-3GalNAc(β-COOH |
| H-type-5-COOH | [100], [500], [1000] | Fucα1-2Galβ1-3Gal(β-COOH |
| 2'FL | [950] | Fucα1-2Galβ1-4Glc(β-Sp |
| H-type-6-COOH | [100], [500], [1000] | Fucα1-2Galβ1-4Glc(β-COOH |
| A tetra type 1 | [700] | GalNAcα1-3[Fucα1-2]Galβ1-3GlcNAc(β-Sp |
| A-type-1-COOH | [135], [540], [1350] | GalNAcα1-3[Fucα1-2]Galβ1-3GlcNAc(β-COOH |
| A tetra type 2 | [920] | GalNAcα1-3[Fucα1-2]Galβ1-4GlcNAc(β-Sp |
| A-type-2-COOH | [135], [540], [1350] | GalNAcα1-3[Fucα1-2]Galβ1-4GlcNAc(β-COOH |
| A-type-3-COOH | [135], [540], [1350] | GalNAcα1-3[Fucα1-2]Galβ1-3GalNAc(α-COOH |
| A-type-4-COOH | [100], [500], [1000] | GalNAcα1-3[Fucα1-2]Galβ1-3GalNAc(β-COOH |
| A-type-5-COOH | [135], [540], [1350] | GalNAcα1-3[Fucα1-2]Galβ1-4Gal(β-COOH |
| A tetra type 6 | [590] | GalNAcα1-3[Fucα1-2]Galβ1-4Glc(β-Sp |
| A-type-6-COOH | [135], [540], [1350] | GalNAcα1-3[Fucα1-2]Galβ1-4Glc(β-COOH |
| B tetra type 1 | B tetra type 1-[620] | Galα1-3[Fucα1-2]Galβ1-3GlcNAc(β-Sp |
| B-type-1-COOH | [135], [540], [1350] | Galα1-3[Fucα1-2]Galβ1-3GlcNAc(β-COOH |

| | | |
|---|---|---|
| B tetra type 2 | [970] | Galα1-3[Fucα1-2]Galβ1-4GlcNAc(β-Sp |
| B-type-2-COOH | [540] | Galα1-3[Fucα1-2]Galβ1-4GlcNAc(β-COOH |
| B-type-3-COOH | [100], [500], [1000] | Galα1-3[Fucα1-2]Galβ1-3GalNAc(α-COOH |
| B-type-4-COOH | [135], [540], [1350] | Galα1-3[Fucα1-2]Galβ1-3GalNAc(β-COOH |
| B-type-5-COOH | [540], [1350] | Galα1-3[Fucα1-2]Galβ1-3Gal(β-COOH |
| B tetra type 6 | [920] | Galα1-3[Fucα1-2]Galβ1-4Glc(β-Sp |
| B-type-6-COOH | [135], [540], [1350] | Galα1-3[Fucα1-2]Galβ1-4Glc(β-COOH |
| 2'F-B type 2 | [220], [760] | Galα1-3[Fucα1-2]Galβ1-4[Fucα1-3]GlcNAc(β-Sp |
| H2 | [430] | Fucα1-2Galβ1-4GlcNAcβ1-3Galβ1-4GlcNAc(β-Sp |
| H3 | [190] | Fucα1-2Galβ1-4GlcNAcβ1-3Galβ1-4GlcNAcβ1-3Galβ1-4GlcNAc(β-Sp |
| Tri-AN3 | [1080] | GalNAcα1-3[Fucα1-2]Gal(β-Sp |
| LeA | [950] | Galβ1-3[Fucα1-4]GlcNAc(β-Sp |
| Lex | [810] | Galβ1-4[Fucα1-3]GlcNAc(β-Sp |
| Gala3Lex | [620] | Galα1-3Galβ1-4[Fucα1-3]GlcNAc(β-Sp |
| Di-Lex | [410] | Galβ1-4[Fucα1-3]GlcNAcβ1-3Galβ1-4[Fucα1-3]GlcNAcβ1-Sp |
| Ley-Lex | [350] | Fucα1-2Galβ1-4[Fucα1-3]GlcNAcβ1-3Galβ1-4[Fucα1-3]GlcNAc(β-Sp |
| LeALex | [350] | Galβ1-3[Fucα1-4]GlcNAcβ1-3Galβ1-4[Fucα1-3]GlcNAc(β-Sp |
| Lex-LeA | [410] | Galβ1-4[Fucα1-3]GlcNAcβ1-3Galβ1-3[Fucα1-4]GlcNAc(β-Sp |
| Lec-LeX | [570] | Galβ1-3GlcNAcβ1-3Galβ1-4[Fucα1-3]GlcNAc(β-Sp |
| Tri-Lex | [430] | Galβ1-4[Fucα1-3]GlcNAcβ1-3Galβ1-4[Fucα1-3]GlcNAcβ1-3Galβ1-4 [Fucα1-3]GlcNAc(β-Sp |

| | | |
|---|---|---|
| Ley-Di-Lex | [510] | Fucα1-2Galβ1-4[Fucα1-3]GlcNAcβ1-3Galβ1-4[Fucα1-3]GlcNAcβ1-3 Galβ1-4[Fucα1-3]GlcNAc(β-Sp |
| aMan | [840] | Man(α-S6 |
| (Man)3 | [<8] | Manα1-6[Manα1-3]Man(α-S6 |
| | [1300] | Manα1-6[Manα1-3]Man(α-S6 |
| | [1730] | Manα1-6[Manα1-3]Man(α-S6 |
| 11 | [50], [150], [500], [750], [780] | Manα1-6[Manα1-3]Manβ1-4GlcNAcβ1-4GlcNAc(β1-Sp |
| 6 | [50], [150], [500], [750], [810], [1000] | GlcNAcβ1-2Manα1-6[GlcNAcβ1-2Manα1-3]Manβ1-4GlcNAcβ1-4GlcNAc(β1-Sp |
| 10 | [150], [500], [540], [1000] | Galβ1-4GlcNAcβ1-2Manα1-6[Galβ1-4GlcNAcβ1-2Manα1-3]Manβ1-4GlcNAcβ1-4GlcNAc(β1-Sp |
| 9 | [50], [510], [730], [950], [970] | Neu5Acα2-6Galβ1-4GlcNAcβ1-2Manα1-6[Neu5Acα2-6Galβ1-4GlcNAcβ1-2 Manα1-3]Manβ1-4GlcNAcβ1-4GlcNAc(β1-Sp |
| GM1 | [460] | Neu5Acα2-3[Galβ1-3GalNAcβ1-4]Galβ1-4Glc(β-Sp |
| GM2 | [190] | Neu5Acα2-3[GalNAcβ1-4]Galβ1-4Glc(β-Sp |
| CT\|Sda | [350] | Neu5Acα2-3[GalNAcβ1-4]Galβ1-4GlcNAc(β-Sp |
| GD1a | [110] | Neu5Acα2-3[Neu5Acα2-3Galβ1-3GalNAcβ1-4]Galβ1-4Glc(β-Sp |
| 3'S-Di-LeA | [160] | Neu5Acα2-3Galβ1-3[Fucα1-4]GlcNAcβ1-32(β-Sp |
| 3'SLeA-Lex | [570] | Neu5Acα2-3Galβ1-3[Fucα1-4]GlcNAcβ1-3Galβ1-4[Fucα1-3]GlcNAc(β-Sp |
| 3'S-Tri-LeX | [160] | Neu5Acα2-3Galβ1-4[Fucα1-3]GlcNAcβ1-3Galβ1-4[Fucα1-3] GlcNAcβ1-3Galβ1-4[Fucα1-3]GlcNAc(β-Sp |
| 3'Slec | [430] | Neu5Acα2-3Galβ1-3GlcNAc(β-Sp |

| | | |
|---|---|---|
| GM3 | [190] | Neu5Acα2-3Galβ1-4Glc(β-Sp |
| | [540] | Neu5Acα2-3Galβ1-4Glc(β-Sp |
| 3'SLDN | [460] | Neu5Acα2-3GalNAcβ1-4GlcNAc(β-Sp |
| 3'S-Di-Lec | [270] | Neu5Acα2-3Galβ1-3GlcNAcβ1-3Galβ1-3GlcNAc(β-Sp |
| 3'SLecLN | [860] | Neu5Acα2-3Galβ1-3GlcNAcβ1-3Galβ1-4GlcNAc(β-Sp |
| 3'SLN-Lec | [410] | Neu5Acα2-3Galβ1-4GlcNAcβ1-3Galβ1-3GlcNAc(β-Sp |
| 3'S-Di-LN | [1350] | Neu5Acα2-3Galβ1-4GlcNAcβ1-3Galβ1-4GlcNAcβ1-Sp |
| 3'STri-LN | [160] | Neu5Acα2-3Galβ1-4GlcNAcβ1-3Galβ1-4GlcNAcβ1-3Galβ1-4GlcNAc(β-Sp |
| 3'SLec (Gc) | [350] | Neu5Gcα2-3Galβ1-3GlcNAc(β-Sp |
| 3'SL (Gc) | [490] | Neu5Gcα2-3Galβ1-4Glc(β-Sp |
| 3'SLN (Gc) | [570] | Neu5Gcα2-3Galβ1-4GlcNAc(β-Sp |
| 3'-KDNLec | [760] | Kdnα2-3Galβ1-3GlcNAc(β-Sp |
| 3'KDNLN | [510] | Kdnα2-3Galβ1-4GlcNAc(β-Sp |
| 6'SL | [620] | Neu5Acα2-6Galβ1-4Glc(β-Sp |
| 6'SLN (Gc) | [430] | Neu5Gcα2-6Galβ1-4GlcNAc(β-Sp |
| 6'SLDN | [620] | Neu5Acα2-6GalNAcβ1-4GlcNAc(β-Sp |
| 6'SLN-Lec | [300] | Neu5Acα2-6Galβ1-4GlcNAcβ1-3Galβ1-3GlcNAc(β-Sp |
| 6'S-Di-LN | [300] | Neu5Acα2-6Galβ1-4GlcNAcβ1-3Galβ1-4GlcNAcβ1-Sp |
| GD2 | [460] | Neu5Acα2-8Neu5Acα2-3[GalNAcβ1-4]Galβ1-4Glc(β-Sp |
| GD3 | [320] | Neu5Acα2-8Neu5Acα2-3Galβ1-4Glc(β-Sp |
| TetraSLac | [80] | Neu5Acα2-8Neu5Acα2-8Neu5Acα2-8Neu5Acα2-3Galβ1-4Glc(β-Sp |
| GT2 | [160] | Neu5Acα2-8Neu5Acα2-8Neu5Acα2-3[GalNAcβ1-4]Galβ1-4Glc(β-Sp |

| | | |
|---|---|---|
| GT3 | [510] | Neu5Acα2-8Neu5Acα2-8Neu5Acα2-3Galβ1-4Glc(β-Sp |
| GQ2 | [140] | Neu5Acα2-8Neu5Acα2-8Neu5Acα2-8Neu5Acα2-3[GalNAcβ1-4]Galβ1-4Glc(β-Sp |
| (Galf)4 | [<8] | Gal*f*β1-5Gal*f*β1-5Gal*f*β1-5Gal*f*(β-S8 |
| L-Glca-C8 | [25], [50], [150], [500], [4000] | L-Glcα-O-octyl-COOH |
| L-Glcb-C8 | [25], [50], [150], [500], [4000] | L-Glcβ-O-octyl-COOH |
| Glcb-C4 | [25], [50], [150], [500], [4000] | D-Glcβ-O-butyl-COOH |
| L-Gala-C8 | [25], [50], [150], [500], [4000] | L-Galα-O-octyl-COOH |
| L-Galb-C8 | [25], [50], [150], [500], [4000] | L-Galβ-O-octyl-COOH |
| Galb-C4 | [25], [50], [150], [500], [4000] | D-Galβ-O-butyl-COOH |
| L-Mana-C8 | [25], [50], [150], [500], [4000] | L-Manα-O-octyl-COOH |
| Mana-C4 | [25], [50], [150], [500], [4000] | D-Manα-O-butyl-COOH |
| L-Fuca-C8 | [25], [50], [150], [500], [4000] | L-Fucα-O-octyl-COOH |
| L-Fucb-C8 | [25], [50], [150], [500], [4000] | L-Fucβ-O-octyl-COOH |
| D-Fuca-C8 | [25], [50], [150], [500], [4000] | D-Fucα-O-octyl-COOH |
| D-Fucb-C8 | [25], [50], [150], [500], [4000] | D-Fucβ-O-octyl-COOH |
| AzOH | [460] | 2-Azidoethanol |
| | [500] | 2-Azidoethanol |
| | [950] | 2-Azidoethanol |
| blank | [0] | Wild-type phage (no modification) |

**Table S1. List of glycans in LiGA-ED and LiGA-NOA.**
Glycans that are presented in LiGA were highlighted in green.

| Strain | Host Origin | Source |
|---|---|---|
| *L. reuteri* DSM20016. Human | Human | JGI 2671180761 |
| *L. reuteri* mm2-3. Human | Human | JGI 2502171171 |
| *L. reuteri* SD2112.human | Human | JGI 650716048 |
| *L. reuteri* M27U15.human | Human | JGI 2687453659 |
| *L. reuteri* i5007.pig | Porcine | JGI 2554235423 |
| *L. reuteri* lp167. Pig | Porcine | JGI 2599185361 |
| *L. reuteri* ATCC53608. Pig | Porcine | EMBL LN906634 |
| *L. reuteri* limo.pig | Porcine | Willing's lab |
| *L. reuteri* JCM 1081.poultry | Poultry | JGI 2684623011 |
| *L. reuteri* 1366.poultry | Poultry | JGI 2684623010 |
| *L. reuteri* AP3.Poultry | Poultry | GCA_014145445 |
| *L. reuteri* CSF8.poultry | Poultry | JGI 2684623009 |
| *L. reuteri* MLC3.mouse | Rodent | JGI 2506381016 |
| *L. reuteri* TMW1.656. rat | Rodent | JGI 2534682350 |
| *L. reuteri* LPUPH1.rat | Rodent | JGI 2506381017 |
| *L. reuteri* 100-23. rat | Rodent | JGI 2500069000 |

**Table S2. List of L. reuteri strains used in this study.**

| **Bacteria** | **Growth media** | **Host** |
| --- | --- | --- |
| *B. vulgatus* | BHI + 0.5 L-cysteine + 5% yeast extract | Human |
| *B. fragilis* | BHI + 0.5 L-cysteine + 5% yeast extract | Human |
| *B. dorei* | BHI + 0.5 L-cysteine + 5% yeast extract | Human |
| Mice *E. coli* (commensal) | LB | Mice |
| Rat *E. coli* (commensal) | LB | Rat |
| *Clotridium ramosum* | BHI + 0.5 L-cysteine/ FAA | Human |
| *Citrobacter freundii* | BHI + 0.5 L-cysteine/ FAA | Human |
| *AIEC* | LB | Human |
| *ETEC* | LB | Human |
| *Bacteroides thetaiotamicron* | BHI + 0.5 L-cysteine + yeast extract | Porcine |
| *Bacteroides thetaiotamicron* | BHI+0.5 L-cysteine + yeast extract | Human |
| *Lactobacillus mucosae* | MRS + 5% maltose + 10% fructose | Porcine |

**Table S3. Bacterial strains used in this study, including growth media and host origin.**
BHI, brain heart infusion medium; LB, Luria-Bertani medium; FAA, fastidious anaerobe agar; MRS, de Man, Rogosa, and Sharpe medium.

| **Bacteria** | **CFU $ml^{-1}$ of broth** | **OD600** | **Incubation time (growth time)** |
|---|---|---|---|
| *L. reuteri* | $1.6 \times 10^9$ | 1.11 | 18 h |
| *E. coli* | $1.8 \times 10^9$ | 1.10 | 18 h |
| *B. vulgatus* | $3 \times 10^8$ | 1.00 | 48 h |
| *B. fragilis* | $6 \times 10^8$ | 1.00 | 48 h |
| *B. dorei* | $5\times 10^8$ | 1.20 | 48 h |
| Mice *E. coli* (commensal) | $1.8 \times 10^9$ | 1.10 | 18 h |
| Rat *E. coli* (commensal) | $1.8 \times 10^9$ | 1.10 | 18 h |
| *Clotridium ramosum* | $1.6 \times 10^8$ | 1.07 | 18 h |
| *Citrobacter freundii* | $5 \times 10^8$ | 1.09 | 18 h |
| *AIEC* (*E. coli* from Human) | $1.8 \times 10^9$ | 1.10 | 18 h |
| *ETEC* (*E. coli* from Human) | $1.8 \times 10^9$ | 1.10 | 18 h |
| *Bacteroides thetaiotamicron* | $1.8 \times 10^9$ | 1.08 | 24 h |
| *Lactobacillus mucosae* | $1.8 \times 10^9$ | 1.09 | 18 h |

**Table S4. CFU and OD600 enumeration.**
CFU, colony-forming unit; OD600, optical density at 600 nm.

| LiGA | Dataset* | Target | Comment |
|---|---|---|---|
| LiGA-ED | 20210813-87EDbwpnGT-OB | w.t. *E. coli* BW25113 | Used in **Fig. 1D–F** |
| | 20210813-87EDbwnnGT-OB | ΔFimH *E. coli* BW25113 | |
| | 20210813-87EDnsdGT-OB | No target (LiGA input) | |
| | 20211004-87EDnsdGT-OB | | |
| | 20211115-87EDnsdGT-OB | | |
| | 20211216-87EDnsdGT-OB | | |
| | 20210813-87EDjcmGT-OB | *L. reuteri* JCM1081 (Poultry) | Used in **Fig. 2–3** and **Fig. S1–S4** |
| | 20211004-87EDjcmGT-OB | *L. reuteri* JCM1081 (Poultry) | |
| | 20211115-87EDjcmGT-OB | *L. reuteri* JCM1081 (Poultry) | |
| | 20211115-87EDlimrGT-OB | *L. reuteri* limo (Porcine) | |
| | 20210813-87EDrapGT-OB | *L. reuteri* AP3 (Poultry) | |
| | 20210813-87EDmmGT-OB | *L. reuteri* mm2-3 (Human) | |
| | 20210813-87EDlratGT-OB | *L. reuteri* ATCC53608 (Porcine) | |
| | 20210813-87EDcsfGT-OB | *L. reuteri* CSF8 (Poultry) | |
| | 20210813-87EDmlGT-OB | *L. reuteri* MLC3 (Rodent) | |
| | 20210813-87EDsdlrGT-OB | *L. reuteri* SD2112 (Human) | |
| | 20210813-87EDfaaGT-OB | *L. reuteri* 1366 (Poultry) | |
| | 20210813-87EDgaaGT-OB | *L. reuteri* i5007 (Porcine) | |
| | 20210813-87EDdsmGT-OB | *L. reuteri* DSM 20016 (Human) | |
| | 20210813-87EDlplrGT-OB | *L. reuteri* lp167 (Porcine) | |
| | 20210813-87EDlpuphGT-OB | *L. reuteri* LPUPH1 (Rodent) | |
| | 20210813-87EDrodGT-OB | *L. reuteri* 100-23 (Rodent) | |
| | 20210813-87EDtmwGT-OB | *L. reuteri* TMW1.656 (Rodent) | |
| | 20210813-87EDdaaGT-OB | *L. reuteri* M27U15 (Human) | |
| | 20211115-87EDcfhGT-OB | *C. fruendii* (Human) | Used in **Fig. 3** and **Fig. S5–S6** |
| | 20211216-87EDcfhGT-OB | *C. fruendii* (Human) | |
| | 20211115-87EDcrhGT-OB | *C. ramosum* (Human) | |
| | 20211216-87EDcrhGT-OB | *C. ramosum* (Human) | |
| | 20211115-87EDlmucGT-OB | *L. mucosae* (Porcine) | |
| | 20211115-87EDbfraGT-OB | *B. fragilis* (Human) | |
| | 20211115-87EDbtpGT-OB | *B. thetaiotamicron* (Porcine) | |
| | 20211216-87EDbtpGT-OB | *B. thetaiotamicron* (Porcine) | |
| | 20211115-87EDbvulGT-OB | *B. vulgatus* (Human) | |
| | 20211216-87EDbvulGT-OB | *B. vulgatus* (Human) | |
| | 20211115-87EDbtheGT-OB | *B. thetaiotamicron* (Human) | |
| | 20211216-87EDbtheGT-OB | *B. thetaiotamicron* (Human) | |
| | 20211115-87EDbdorGT-OB | *B. dorei* (Human) | |
| | 20211115-87EDrateGT-OB | Rat *E. coli* (commensal) | Used in **Fig. 3** and **Fig. 4** |
| | 20211216-87EDrateGT-OB | Rat *E. coli* (commensal) | |
| | 20211115-87EDmcenGT-OB | Mice *E. coli* (commensal) | |

| | | | |
|---|---|---|---|
| | 20211216-87EDmcenGT-OB | Mice *E. coli* (commensal) | |
| | 20211115-87EDaiecGT-OB | *AIEC* (*E. coli* from Human) | |
| | 20211115-87EDetecGT-OB | *ETEC* (*E. coli* from Human) | |
| LiGA-NOA | 20251218-87NOAoraCAAA-EG | *L. reuteri* JCM 1081 | Used in **Fig. 5A** |
| | 20251218-87NOAooCAAA-EG | No target (LiGA input) | |
| | 20251218-87NOAepaDC-IF | w.t. *E. coli* BW25113 | Used in **Fig. 5B–C** |
| | 20251218-87NOAeqaDC-IF | ΔFimH *E. coli* BW25113 | |
| | 20251218-87NOAooNA-IF | No target (LiGA input) | |

**Table S5. LiGA data used in this work.**
The listed datasets are publicly available at https://48hd.cloud/.

### References

(1) Coolen, M. J.; Post, E.; Davis, C. C.; Forney, L. J. Characterization of microbial communities found in the human vagina by analysis of terminal restriction fragment length polymorphisms of 16S rRNA genes. *Appl. Environ. Microbiol.* **2005**, *71* (12), 8729–8737.

(2) Sojitra, M.; Sarkar, S.; Maghera, J.; Rodrigues, E.; Carpenter, E. J.; Seth, S.; Ferrer Vinals, D.; Bennett, N. J.; Reddy, R.; Khalil, A. Genetically encoded multivalent liquid glycan array displayed on M13 bacteriophage. *Nat. Chem. Biol.* **2021**, *17* (7), 806–816.

(3) Lin, C.-L.; Sojitra, M.; Carpenter, E. J.; Hayhoe, E. S.; Sarkar, S.; Volker, E. A.; Wang, C.; Bui, D. T.; Yang, L.; Klassen, J. S. Chemoenzymatic synthesis of genetically-encoded multivalent liquid N-glycan arrays. *Nat. Commun.* **2023**, *14* (1), 5237. Reddy, R.; Carpenter, E.; Halpin, A.; Sojitra, M.; Peng, C.; Lima, G. M.; Xue, X.; Yan, K.; Pearcy, J.; Ellis, M. Evaluation of Multiplexed Liquid Glycan Array (LiGA) for Serological Detection of Glycanbinding Antibodies. *Glycobiology* **2025**, cwaf042. Carpenter, E. J.; Peng, C.; Haregu, S.; Twells, N.; Woudstra, L.; Sood, A.; Cartmell, J.; Woods, R. J.; Mahal, L. K.; Wang, S.-K. Atom-level machine learning of protein-glycan interactions and cross-chiral recognition in glycobiology. *Sci. Adv.* **2025**, *11* (49), eadx6373.

(4) Sojitra, M.; Schmidt, E. N.; Lima, G. M.; Carpenter, E. J.; McCord, K. A.; Atrazhev, A.; Macauley, M. S.; Derda, R. Measuring carbohydrate recognition profile of lectins on live cells using liquid glycan array (LiGA). *Nat. Protoc.* **2025**, *20* (4), 989–1019.

(5) Matochko, W. L.; Cory Li, S.; Tang, S. K.; Derda, R. Prospective identification of parasitic sequences in phage display screens. *Nucleic Acids Res.* **2014**, *42* (3), 1784–1798.

(6) Robinson, M. D.; Smyth, G. K. Small-sample estimation of negative binomial dispersion, with applications to SAGE data. *Biostatistics* **2008**, *9* (2), 321–332.

(7) Benjamini, Y.; Hochberg, Y. Controlling the false discovery rate: a practical and powerful approach to multiple testing. *J. R. Stat. Soc. Ser. B* **1995**, *57* (1), 289–300.

(8) Robinson, M. D.; Oshlack, A. A scaling normalization method for differential expression analysis of RNA-seq data. *Genome Biol.* **2010**, *11* (3), R25.
